# Lineage-Specific Evolution, Structural Diversity, and Activity of R2 Retrotransposons in Animals

**DOI:** 10.1101/2025.05.05.652312

**Authors:** Nozhat T. Hassan, Briana Van Treeck, Anthony Rodríguez-Vargas, Anna E. Sheppard, Kathleen Collins, David L. Adelson

## Abstract

Retrotransposons play outsized roles in the evolution of gene regulation, genome function, and disease pathogenesis and more recently, have sparked interest as workhorses for new gene therapy approaches. R2 retrotransposons insert site-specifically to the multicopy genes encoding 28S ribosomal RNA at a target sequence conserved broadly across eukaryotic evolution. R2 retrotransposons have been detected in many animals, but previous surveys have been limited in scope and methodology. Here, we substantially expand the known distribution of R2 retrotransposons from previously unrepresented or underrepresented taxonomic groups ranging from ctenophores to amphibians and reptiles. We discover diverse R2 domain architectures and motifs and identify many new avian R2 candidates for genome engineering development. Overall, phylogenetic analyses reveal two highly successful R2 lineages. We observe properties of each lineage in several features of the domains that mediate DNA recognition and in co-evolving signatures within the reverse transcriptase domain. Within each of the two lineages, R2 protein sequences do not necessarily preserve the unifying configuration of N-terminal DNA-binding domains implied in the current clade classification scheme. We show that recombinant R2 proteins with distinctive domain architectures and distribution across major animal classes support target-primed reverse transcription with conserved site specificity. Our analysis of the surprisingly varied domain architectures that support target-site specificity informs new R2 classification criteria and provides a greatly expanded foundation for additional structure/function insights about DNA binding selectivity. This expanded perspective on R2 evolution informs approaches for engineering therapeutic gene insertion technologies and offers an opportunity to investigate the conservation and diversification of retrotransposons.

## Introduction

Non-long terminal repeat (non-LTR) retrotransposons are generally ubiquitous across eukaryotes, contributing to both genome evolution and disease through their mobility and other activities *(1-4)*. Non-LTR retrotransposons are present across animals in different combinations of retrotransposon identities and abundances, suggestive of dramatically differential mobility and molecular evolution resulting from arms races with transposon repression mechanisms within the host genome *(5)*. Recent efforts have adapted non-LTR retrotransposon machinery into new tools for gene insertion, promising potential new approaches for individualised disease therapy. Non-LTR retrotransposons with minimal target-site sequence requirements can be used to randomly place gene insertions or can be fused with targeting determinants *(6, 7)*. On the other hand, retrotransposons that have evolved very restrictive target-site specificity are useful for gene addition to specific loci, preferably to safe harbour regions of the human genome *(8)*. Precise RNA-mediated insertion of transgenes (PRINT) is one such technology that relies on the site-specificity of one of the evolutionarily most widespread non-LTR retrotransposons, R2 *(9)*.

The R2 family of non-LTR retrotransposons inserts site-specifically into the 28S ribosomal RNA (rRNA) region of genes encoding the large rRNA precursor (rDNA loci), present in genomes as tandemly repeated rDNA arrays *(10)*. R2s lack an internal promoter and at least in some species, are dependent on the rDNA RNA polymerase I promoter for proliferation *(10, 11)*. The single R2-encoded protein (R2p) recognises and nicks the 28S target site and uses the nicked DNA as a primer for cDNA synthesis directly into the genome, a process termed target-primed reverse transcription (TPRT) *(10, 12)*. This requires coordinated R2p reverse transcriptase (RT) and endonuclease (EN) activities *(12, 13)*. The most N-terminal portion of R2p contains the majority of the DNA-binding surface for the target site *(14-16)*. Different domain architectures of the N-terminal region, with a variable 1-3 zinc-fingers (ZnFs) preceding an adjacent Myb domain, have been the basis of classifying R2 elements into 4 clades further divided into 11 subclades *(9, 17-19)*. ZnFs are numbered with the conserved ZnF1 sitting N-terminal to the Myb domain and subsequent ZnFs progressing towards the N-terminus, with ZnF2 differing from ZnF1 and ZnF3 in a cysteine (C) and histidine (H) pattern of CCHC instead of CCHH. Current R2 classification thus has clade A (ZnF3+ZnF2+ZnF1), clades B and C (ZnF2+ZnF1 and ZnF3+ZnF1, respectively), and clade D (ZnF1 only), with A-clade and D-clade R2p appearing most common *(19)*.

R2s were discovered within the genome of *Drosophila melanogaster* (fruit fly) and were thought to be intronic sequences, but this was disputed when genes with R2 insertions did not appear to be transcribed *(20)*. R2s have since been found in a variety of organisms through the generation of genomic sequence data. They are notably found in Ctenophora and extend to Reptilia and Aves but not mammals *(19)*. This indicates that R2s have persisted since near the beginning of multicellular eukaryotic life approximately 800 million years ago *(21)*. However, too few sequences have been identified to date to understand the phylogenetic scope of full-length R2 persistence and whether particular R2 N-terminal domain configurations occur only in specific phylogenetic groups. Prior to this work, the known distribution of R2s from RepBase *(22-24)* consisted of 185 sequences from 144 species, with the majority from arthropods; just under 60 were full-length R2s with an entire, unambiguous R2p open reading frame (ORF) with flanking 5’ and 3’ untranslated regions (UTRs) embedded in 28S rDNA.

Here, we created an R2 discovery workflow that used long-read sequencing data and genome assemblies to curate over 330 new R2s from major classes of Metazoan organisms. We identified the first R2s in Amphibia and Chondrichthyes and significantly expanded known R2s in reptiles, particularly in snakes. We discovered a surprising diversity of R2 N-terminal domain structures that does not fall within the current classification scheme *(17, 19)*. This diversification is overlaid on the propagation of two anciently diverged and highly successful lineages, each with co-evolving domains, one lineage with a co-folded zinc finger pair of nucleic acid interaction domains that appears to be unique to R2s. Our biochemical and cellular assays of a broad sampling of recombinant R2p expressed in human cells confirm the conservation of target site specificity despite large relative differences in endonuclease and TPRT activities. This thorough expansion of R2 phylogeny annotation and functional testing enables deeper insights into retrotransposon evolution as well as R2p structure/function and engineering.

## Results

### Significant expansion of R2 retrotransposons across Metazoa

We aimed to sample R2 retrotransposon phylogeny more broadly than previously undertaken to gain a deeper insight into how R2s have diversified during Metazoan evolution. Many sequences of R2s reported to date are incomplete, with only a partial ORF or truncated UTRs *(22-24)*. Furthermore, because genome sequencing in the past largely relied on short-read sequencing, R2 consensus sequences derived from alignments of multiple reads, for example, in the most recently expanded R2 inventory, may not accurately represent an actual full-length R2 ORF *(19)*. With the increasing availability of long-read data and improved genome assemblies, we sought to detect potentially functional R2s more comprehensively across major animal groups.

Initially, our strategy to find R2s in genomes of interest mirrored past studies: we used BLASTN+ to search the genome of interest with R2 query sequences from each previously defined clade (A, B, C, and D). The top hits were extended in both the 5’ and 3’ directions to find flanking 28S rDNA sequences *(19)*. Depending on whether both sides of the target site were located, sequences were annotated as full-length R2s or truncated R2s. However, this strategy often failed to return full-length R2s. Genome assemblies are not necessarily reliable regarding repetitive structures, especially of R2s, which are repeats nested within the larger repeats of an rDNA array *(19, 25)*. We hypothesised that some missing motifs could be due to misassembly rather than the absence of a full-length R2 in the organism. To address this limitation, we parsed long-read genome sequencing data with R2 query sequences when genome assemblies returned truncated R2 open reading frames (ORFs) (SI Table 1).

Overall, after querying 360 genomes, we retrieved sequences from 300 total species in Actinopterygii, Amphibia, Arthropoda, Chondrichthyes, Cnidaria, Ctenophora, Echinodermata, Mollusca, Platyhelminthes, Porifera, Reptilia (Aves, Rhynchocephalia, Squamata, Testudines), and Tunicates (SI Table 2). Our focus was on phylogeny beyond the taxonomic groups most frequently sampled in past R2 studies, thereby revealing a more widespread distribution of R2s than previously recognised (Fig. 1a) *(19, 26)*. This includes the discovery of an expansion of R2s in Chondrichthyes (cartilaginous fish), Actinopterygii (ray-finned fish), and Reptilia, and the first reported amphibian R2s from salamanders (Fig. 1a-b). Full-length R2s in Reptilia persist across the wide phylogenetic diversity of Sphenondon (Tuatara), Squamata (snakes and lizards), Crocodilia, and Testudines (turtles) (Fig. 1a). Previously, the few identified snake and lizard R2s were truncated (19); nonetheless, we recovered full-length R2 from many species in these groups.

**Figure 1:**
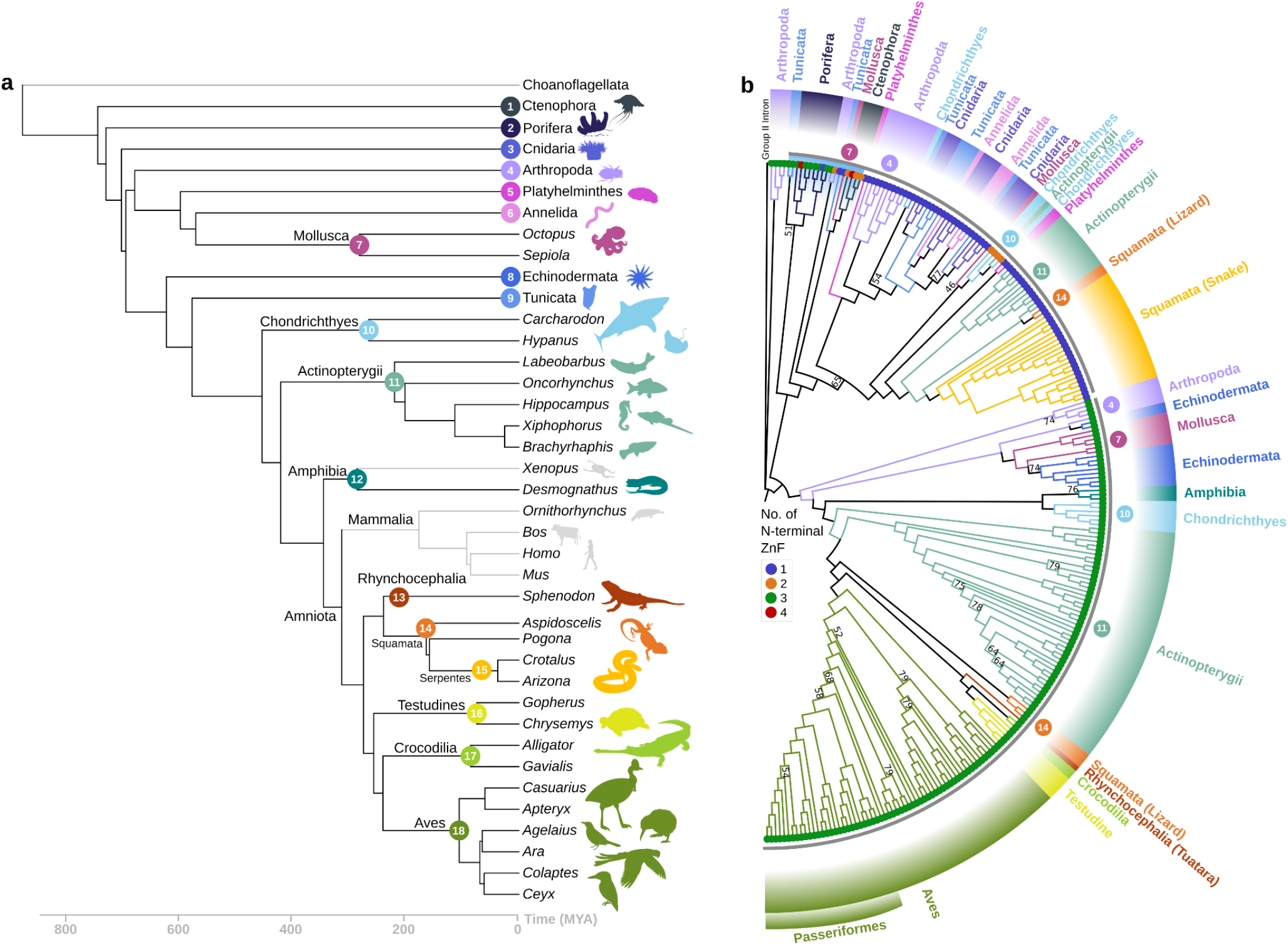
R2 retrotransposons are widely distributed across Metazoa but do not conform to the host species phylogeny. (a) Simplified divergence of major classes of animals relevant to this work, with Ctenophora as the sister group to all other animals, made with TimeTree (58). Time is shown as millions of years ago (MYA) on the bottom axis. Groups with no known R2s are shown in grey. The coloured numbers at some nodes of the host tree are used to number major phyla that are used again in (b). (b) Phylogenetic tree of R2 ORF amino acid sequences. Only tree topology is shown. Branch lengths displayed are arbitrary, only the relationships between the sequences are considered. Each tip is appended with a coloured circle symbol (in blue - 1 ZnF, orange - 2 ZnF, green - 3 ZnF, and red - 4 ZnF), designating the number of ZnFs in the N-terminus of the R2p sequence. Working outwards: a small group of early-branching R2s with diverse ZnF arrangements at the top of the panel are highlighted with a light blue arch, and on the same arc trajectory, the grey arcs show the two major R2 lineages. Outer arcs denote phylogenetic groups that the R2s were curated from, with the colour code from (a). Coloured numbers at branches indicate instances where there is a major discordance from the host tree in (a). The order Passeriformes in Aves is annotated. Group II intron from *T. vestitus* was used as the outgroup. IQTree was used for tree building with 1000 replicates, and branch support values under 80 are shown at the nodes (35).

While failure to detect R2 is not an absolute indicator of their absence, we suggest that the lack of identifiable R2 in several phylogenetic groups indicates stochastic evolutionary loss (Fig. 1a, tree lines and organism schematics in light grey). We also did not detect R2s in unicellular organisms, including Choanoflagellates (SI Table 3). As in previous studies, we defined sequences as R2 based on their homology to other known R2 retrotransposons and their insertion into the conserved 28S rDNA target site. We discovered that R2-homologous retrotransposon sequences in Porifera (sponges) did not insert into the R2 target site. Some of these proteins lacked a Myb domain, and these had the highest sequence homology to Utopia non-LTR retrotransposon proteins (SI Table 4), which are often site-specific for U2 small nuclear RNA (snRNA) genes *(19, 27)*. We classified retrotransposons as Utopia rather than R2 if the predicted protein lacked the hallmark R2 Myb domain (Fig. 2a, SI Fig. 1a, SI Table 4). The subset of R2-homologous Porifera retrotransposon proteins that harbour a predicted Myb domain and therefore are included in Figure 1b have variable flanking sequences that are not the canonical R2 28S rDNA insertion site (Fig. 2b, top branch). We classified these as non-site-specific R2-homologous retrotransposons (SI Tables 4-5).

**Figure 2:**
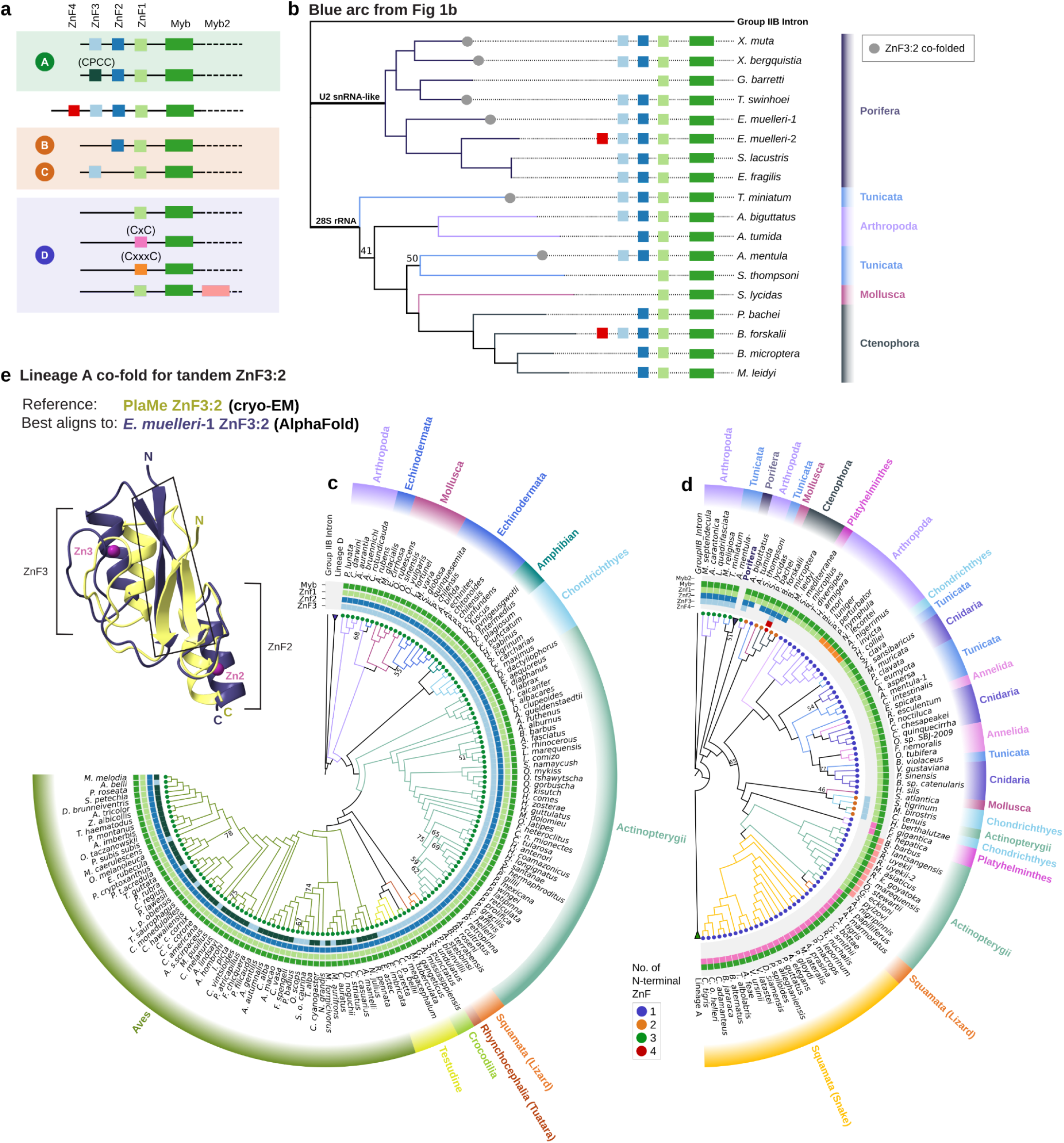
Structural changes in the N terminus of R2p and lineage D and A success. (a) Diversification of ZnF and Myb domain numbers and motif signatures. A - D represents the classification of R2s as clades based on previous studies. ZnF4, ZnF3, ZnF2, and ZnF1 are shown in red, light blue, blue, and light green boxes, respectively. Myb and Myb2 are shown in green and peach boxes. Alternative ZnF3 and ZnF1 motif sequences are annotated as dark green (CxxC to CPCC), pink (CxxC to CxC) and orange (CxxC to CxxxC). (b) Phylogenetic tree of R2s and non-specific R2 elements from the blue arc in Fig. 1b shows the presence or absence of a co-folded ZnF3:2 based on AlphaFold3 predictions. (c,d) Phylogenetic trees with each tip appended with a coloured circle symbol designating the number of zinc fingers in the N-terminus of the R2 amino acid sequence (same as Fig 1b). Coloured boxes for the arc representing each domain match the key in (a). In (c), the tree shows lineage A R2p with exclusively 3 ZnF. In (d), the tree shows lineage D R2p with predominantly D and some A, B and C domain architecture R2s. Porifera R2-homologous elements are collapsed with their domain architecture shown in (b). Arthropod R2s from *M. septendecula, A. carantonica, L. quadrifasciata*, and *M. religiosa* sit outside the major lineage D branch (e) AlphaFold-predicted structure of ZnF3:2 from *E. muelleri*-1 (purple) superimposed with cryo-EM structure of PlaMe (yellow) ZnF3:2 (30). Zinc atoms are shown in magenta. ZnFs in *E. muelleri-1* are all CCHH type, in contrast to PlaMe CCHH and CCHC. Boxed in the aligned structures is the region where most non-covalent, intramolecular interactions concentrate to maintain a single globular unit architecture between ZnFs. IQTree was used for tree building (b-d) with 1000 replicates and branch support values under 80 are shown at the nodes.

### The number of N-terminus ZnF is not a defining characteristic of evolutionary lineage

For additional analysis, we translated our newly curated R2 ORFs. R2 ORFs are non-canonically translated and can lack an initiating methionine *(11, 28)*. We narrowed down our R2 dataset by filtering out R2s with an ORF less than 900 amino acids as this is below the typical length of a clade D R2p and also ensured R2p had at least one potential N-terminal ZnF. This reduced the generated dataset to 180 R2 sequences. We trimmed each ORF to include 10 amino acids N-terminal to the most N-terminal ZnF and used this amino acid sequence for tree-building. Several observations emerged from this analysis.

First, R2 clustering (Fig. 1b) was discordant with species phylogeny overall (Fig. 1a). The R2 sequences fall predominantly into two major branches (Fig. 1b, ring with two grey arcs). Within the 3-ZnF branch (Fig. 1b, lower grey arc), for the most part, R2 phylogeny conforms to host phylogeny. The exception is the 2 groups of Echinodermata R2 split by Mollusca R2s. In the second major branch, there are R2s with 4 to 1 ZnFs (Fig. 1b, upper grey arc), with the majority having 1 ZnF. The early-branching R2 from this group occur in species of Ctenophora, Mollusca, Tunicata, and Arthropoda, and the Porifera R2-homologous retrotransposons fall within this cluster. The Ctenophora R2 from *B. forskalii* encodes a protein with possibly 4 N-terminal ZnFs (Fig. 1b, inner ring red colour, described additionally below) and deviates also in that the flanking target-site rDNA was upstream of the canonical R2 nick site by 11 base-pairs (SI Fig. 2-3). To additionally examine evolutionary diversification, we used other tree-building algorithms rooted with a ctenophore R2p (*B. forskalii)* and obtained largely the same patterns (SI Fig. 4). The exception was a variable placement of the Porifera non-site-specific R2-homologous proteins and their phylogenetically close Tunicata and Arthropoda R2p, which was either the earliest branching of the predominantly 1-ZnF R2p group (Fig. 1b) or the earliest branching of the 3-ZnF R2p group (SI Fig. 4). Ctenophore R2s with 2-4 ZnF consistently clustered with the 1-ZnF group. Additionally, their ZF1-Myb is predicted to make DNA contacts resembling 1-ZnF proteins more than the ZF1-Myb DNA contacts of 3-ZnF group (SI Fig. 1b).

Despite greatly expanding the number and species distribution of R2s, we found only a few with two ZnFs (Fig. 1b, inner ring orange colour). By previous categorisation, these would be designated as B or C clade. Previously, based on phylogenetic clustering, B clade was assigned to R2s in Arthropoda, and the C clade was assigned to R2s in Platyhelminthes, a bitterling fish, and a snake *(10, 19, 29)*. In our expanded R2 dataset, several R2s from Chondrichthyes (sharks and stingrays) with ZnFs that match C clade are clustered with 1 ZnF R2s (Fig. 1b) and not with previously annotated C clade R2s (SI Fig. 5-6) *(10, 19)*. Also surprisingly, most identified Ctenophora R2s have a ZnF configuration that matches the B clade, but they do not cluster with previously annotated B clade R2s (SI Fig. 6-7). The scattered distribution of R2s with 2 ZnF suggests that these domain architectures can arise independently of lineage ancestry rather than as vertically inherited clades of R2.

The distinction between two successful R2 lineages was supported by constructing phylogenetic trees with sequences from different individual domains of R2p: trees based on ZnF1 and Myb, RLE (endonuclease), or Thumb-zinc knuckle domain sequences all resulted in a similar tree topology. Because N-terminal domain architecture does not define phylogenetic proximity, we henceforth refer to the two major phylogenetic branches of R2 as lineages A (Fig. 2c) and D (Fig. 2d) to match the most common number of N-terminus ZnFs to their previous clade A and D designations. We observe a third, more ambiguous branch near the root of the lineage D tree comprising Arthropoda R2s with 3 ZnFs (Fig. 2d: *M. septendecula, A. carantonica, L. quadrifasciata*, and *M. religiosa)*. This branch of R2s appears to cluster with lineage D even when using different protein domains for tree building (SI Fig. 8), but their placement changes to cluster with lineage A when the R2p tree is rooted with Ctenophora R2p (SI Fig. 6.2). Continuing clockwise starting from *T. miniatum* in Figure 2d, additional Arthropoda and some Tunicata R2p group in the lineage D branch with high support values despite structural predictions suggesting that some of these R2p have a co-folded ZnF3:2 (Fig. 2b), a structural property of lineage A R2p (see below). The remaining newly identified unambiguously lineage D R2p include several with domain architectures described previously as being B or C clade (Fig. 2d), here described as B or C domain architecture. Proteins from the previously described B and C clade R2 also cluster in lineage D (SI Fig. 6.1). We note the predominance of 2 ZnF R2s in ctenophore R2p (Fig. 2b) that group with lineage D (Fig. 2d) calls into question the current model that the ancestral R2 was A-clade with 3 ZnF, subsequently giving rise to B, C, and D clades by different ZnF losses *(29)*. Independent of the number of ZnFs in an ancestral R2p, phylogenetic relationships amongst our expanded inventory of R2s suggest that ZnFs can be lost and gained more dynamically than had been anticipated.

### Structural variations in N-terminus ZnFs of lineage D and A R2p

AlphaFold prediction of 3D structures of newly annotated R2p revealed a consistent lineage A feature: interdependently folded ZnF3 and ZnF2 (henceforth, co-folded ZnF3:2) (Fig. 2e). We noticed the presence of this structural feature, although inconsistently, in the non-site-specific R2-homologous Porifera retrotransposon proteins (Fig. 2b, SI Fig. 1c). Nonetheless, when a ZnF3:2 co-fold is predicted like in the case of the *E. muelleri*-1 non-site-specific R2 protein, the fold is superimposable with ZnF3:2 from lineage A R2ps such as from the turtle *P. megacephalum* (Fig. 2e). The *P. megacephalum* R2p ZnF3:2 co-fold has been confirmed by cryo-electron microscopy *(9, 30)*. Intriguingly, we did not find similarly co-folded ZnFs in other proteins by protein structure database searches (See Methods). However, we found that some Utopia non-LTR elements did have similar co-folded ZnFs (ZnF2:1 instead of ZnF3:2 like in R2s) when predicting their N-terminus structure with AlphaFold (SI Figure 14) (See Discussion). The long-branch Arthropoda and Tunicata R2s that cluster near Porifera R2 and share ambiguous tree placement (adjacent to lineage D in Fig. 1b or lineage A in SI Fig. 6.1) either have co-folded ZnF3:2 (the tunicates *T. miniatum* and *A. mentula*, Fig. 2b) or lack it (the arthropod *A. bigittatus*, and 4 additional arthropods; SI Table 6), the latter possibly due to gain of an atypically long linker between ZnF3 and ZnF2. The Ctenophora *B. forskalii* R2p with 4 possible N-terminal ZnFs also lacks the ZnF3:2 co-fold; instead, it has distinctively short linkers between the 3 most N-terminal ZnFs (SI Fig. 1d). These variations complicate the definition of an evolutionary origin for lineage A (see Discussion).

Beyond differences in the number and folding of ZnFs, R2p had additional structural variations in their N-terminal domains (see Fig. 2a colour coding summary). In lineage A R2p, we observed differences in the ZnF3 sequence. Aves R2 ZnF3 often starts with CPCC, which still can form a typical ZnF. However, R2s from some avian species do not have this feature (Fig. 2c, see ZnF3 colour code). Strigiformes (owls) in particular (*P. badius, O. scops, S. o. caurina)* do not have the CPCC ZnF3. Even more structural distinction was evident in lineage D R2p (Fig. 2d; see Fig. 2a colour coding summary). Unusual spacing of cysteines occurred at the start of ZnF1, with CxC instead of CxxC. CxC was detected in snakes, fishes, and flatworms, and CxxxC was detected in arthropods (Fig. 2d, see ZnF1 colour code). Most remarkably, a Myb domain duplication (Fig. 2a, annotated as Myb2) was discovered in a cluster of Actinopterygii fish species from the family Cyprinidae (Fig. 2d). In addition to many ZnF distinctions, lineage D also has more sub-lineages than lineage A, evident in the greater lineage D departure from expectation based on only host organism phylogeny (compare the outer arcs in Fig. 2c-d).

### Different RT motif adaptations in lineage D versus A

We considered how to evaluate R2 diversification by criteria beyond sequence alignments and N-terminal domain architectures. We used position-specific scoring matrices (PSSM) to compare translated RT domains to representative R2p known to be biochemically and biologically active: lineage D R2p RT from the silk moth *B. mori* and lineage A R2p RT from the zebra finch *T. guttata (17)*. PSSM scores (Fig. 3a-d; *B. mori* as BoMo on each y-axis and *T. guttata* as TaGu on each x-axis; see also SI Fig. 9 for detailed annotations) reflect common functionality rather than only sequence identity *(31)*. PSSM score depends on the observation of a particular amino acid in a specific position, and so residues that are both conserved and in a similar position compared to the R2p from *B. mori* or *T. guttat*a receive a higher score. Thus, we would expect a lineage D RT to cluster with BoMo, and *vice versa* with Lineage A RTs with TaGu.

**Figure 3:**
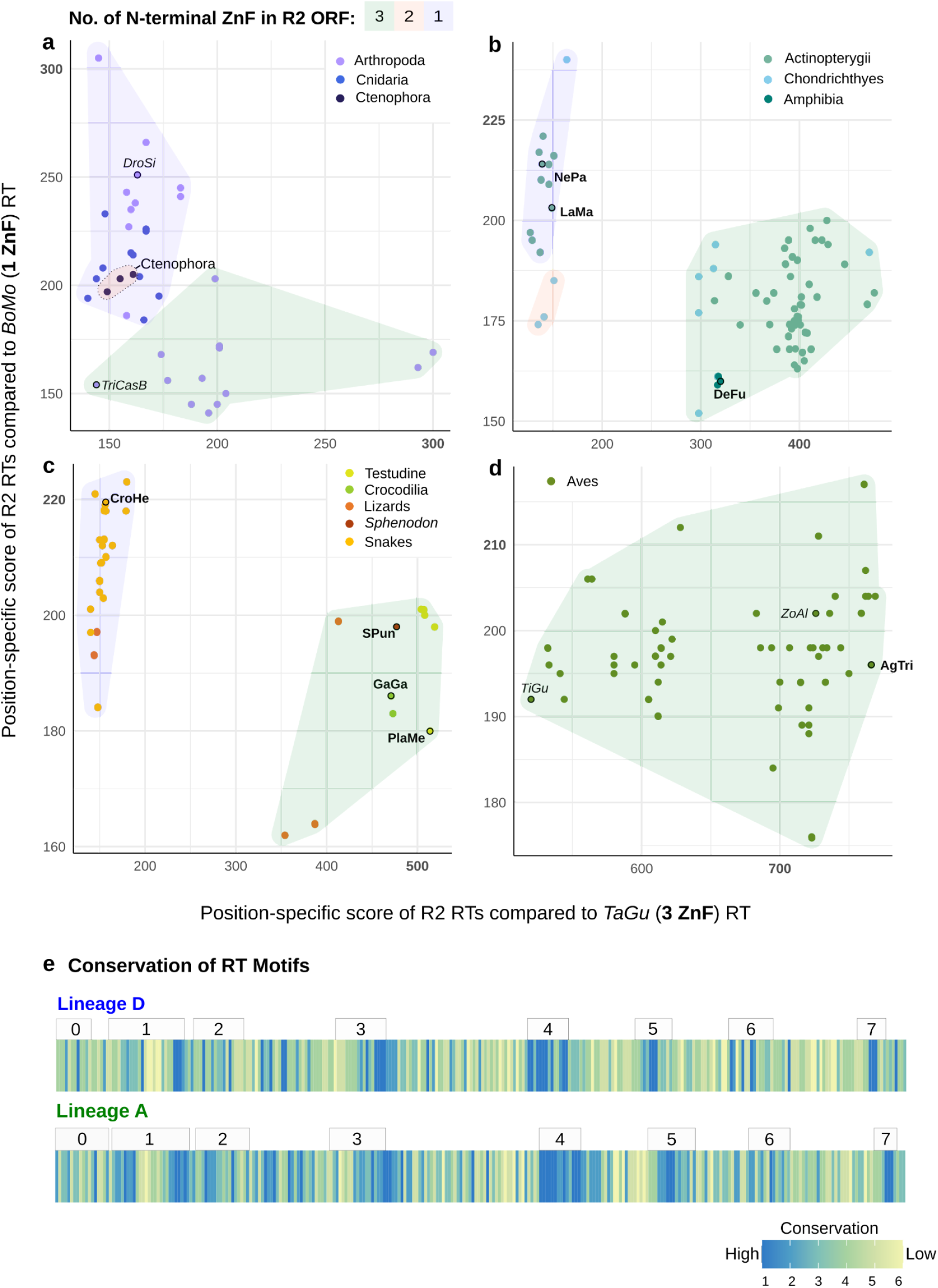
Inferring R2 functionality from RT-domain sequence. PSI-BLAST derived position-specific scoring matrix scores of rDNA-specific R2 RTs compared to BoMo (lineage D) and TaGu (lineage A) RTs. R2s selected for activity testing are circled in black and in bold text. R2s tested in previous studies are in italics: DroSi (*D. simulans)*, TriCasB (*T. castaneum)*, TiGu (*T. guttatus)*, and ZoAl (*Z. albicollis)*. ZoAl R2 sequence was reconstructed from short-reads (See Methods). DroSi, TriCasB, and TiGu are previously characterised sequences (9, 15). Shading in each panel corresponds to the number of N-terminus ZnFs in the complete R2p sequence. R2 RTs from major classes of animals are shown; (a) Arthropoda, Cnidaria, and Ctenophora (b) Amphibia, Actinopterygii, and Chondrichthyes, and (c) Reptilia and (d) Aves. R2s with an atypical number of N-terminus ZnFs (>3) were excluded. (e) Site-specific conservation of individual residues in the RT domain for lineage A and D. Strong conservation of residues is denoted with a lower score (1-3), and rapidly evolving residues are denoted with a high score (4-6). RT motifs from 0 through 7 are annotated.

While only R2p RTs are used in this analysis, we used the number of N-terminus ZnFs to see if R2s still cluster according to the number of N-terminus ZnFs as well as lineage. Comparison of rDNA-specific R2p from early-branching phylogenetic groups (ctenophores, cnidarians, and arthropods) with varying numbers of ZnFs shows that there is as much divergence within a ZnF group as between R2p with different numbers of ZnF (Fig. 3a, see ZnF colour key for background shading and symbol colour key for species). In contrast, amphibian and fish R2p cluster by ZnF (Fig. 3b) as do reptilian R2p (Fig. 3c). Newly identified avian R2p are only from lineage A, as was true for previously reported avian R2p, but two subclusters are observed (Fig. 3d), which are largely delineated by whether the R2 is from a passerine (SI Fig. 9) *(19)*. Passerine R2p include ZoAl from *Z. albicollis* and TaGu, both of which support PRINT transgene insertions in human cells. Non-passerines include R2p from tinamou *T. guttatus*, TiGu, which does not support transgene insertion in human cells *(9)*.

We again used R2p RT domains spanning the well-defined RT motifs from 0-7, to further investigate R2 lineage distinction *(32, 33)*. Using a site-specific rate algorithm, we inferred the evolutionary rate of individual amino acid residues in lineage D or A RT domains (Fig. 3e; SI Fig. 10) *(34, 35)*. A highly conserved site is given a lower score (1-3, blue), whereas a more variable site is given a high score (4-6, yellow). Within lineage A or D, different patterns of sequence conservation were evident (Fig. 3e). The distinct pattern of conserved residues in lineage A compared to lineage D R2p RT domains suggests that RT domain sequence co-evolved with changes in N-terminal domain architecture (see Discussion).

### Co-existence of R2 lineages in phylogenetic groups and individual genomes

In addition to expanding the known phylogenetic breadth of R2 perpetuation, the new R2 inventory also expanded the known co-occurrence of lineage A and lineage D R2s within phylogenetic groups. For example, in addition to arthropods *(18, 19, 32)*, other species also have R2 from both lineage D and A: Squamata (lizards), Actinopterygii, Chondrichthyes, and Mollusca (Fig. 1b, pairs of circled numbers show these instances of lineage A and lineage D phylogenetic co-occurrence). Surprisingly, within these phylogenetic groups, we found several species with full-length R2s from different lineages, as previously described for arthropods *(18, 19, 32)*. These include species of Actinopterygii fish (*L. marequensis, B. barbus)*, Chondrichthyes sharks (*S. tigrinum)*, and Tunicata sea squirts (*A. mentula)* (SI Table 7). Therefore, it is not as rare as we had anticipated that multiple lineages co-exist despite competition for the same target sites.

### Survey of R2p endonuclease and TPRT activities across diverse species

Genome-embedded copies of retrotransposons are often incapable of mobility, which brings into question whether divergent R2p from our expanded phylogeny retain the biochemical activities essential for mobility. We selected newly identified R2p with apparently intact RT motifs, that had different architectures of N-terminal domains to sample for activity, focused on phylogenetic groups beyond the Arthropoda R2s that have been best studied (Fig. 4a, SI Fig. 11). Our final choices included two lineage D fish R2p: LaMa (*L. marequensis*, Largescale yellowfish) with a Myb domain duplication and NePa (*N. papilliferus*, killifish) with CXC ZnF1. We chose lineage A amphibian R2p DeFu (*D. fucus*, Northern dusky salamander) as a representative for the first identification of amphibian R2s, and among reptilian R2s we included a lineage D snake R2p CroHe (*C. o. helleri*, Southern Pacific Rattlesnake) and several lineage A R2p: PBla (*P. blainvillii*, Blainville’s Horned Lizard), SPun (*S. punctatus*, Tuatara), GaGa (*G. gangeticus*, Gharial crocodile), and PlaMe (*P. megacephalum*, Big-headed turtle) subsequently used for structure determination (9, 30). As another avian R2p, we tested AgTri (*A. tricolor*, Tricoloured blackbird).

**Figure 4:**
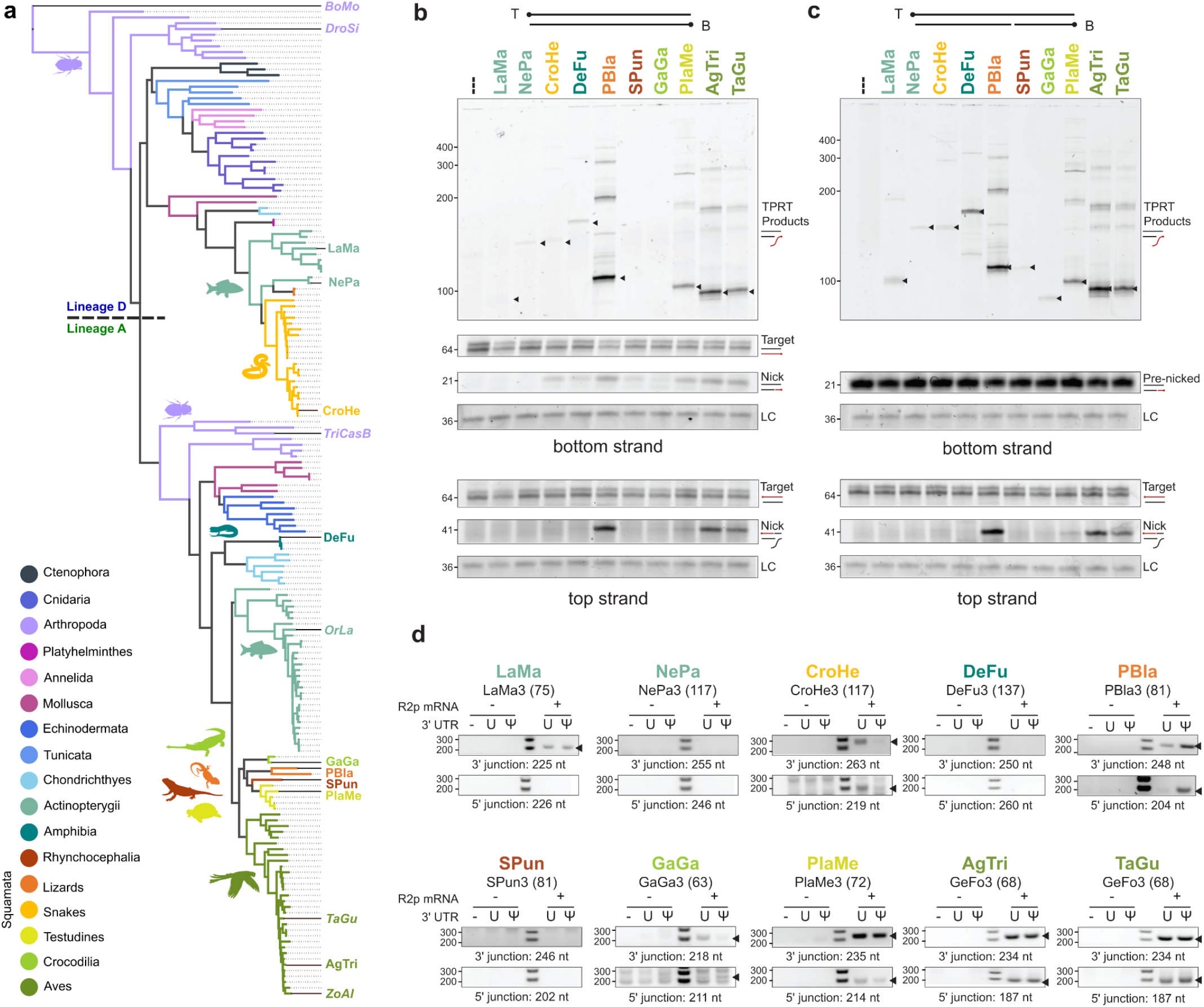
Activity across diverse R2p. (a) Phylogenetic tree of R2 RT domains. R2s annotated at their corresponding branches were selected for R2p activity testing. R2s in italics have been tested in previous studies (9, 15, 70, 71). (b-c) *In vitro* assays using either intact (b) or pre-cleaved (c) target-site oligonucleotide substrates. Relative protein concentration is shown by immunoblot (SI Fig. 13). Expected TPRT product sizes for initial cDNA synthesis products are denoted with a black triangle; note that longer products can form by template-jumping, due to excess RNA in the reactions (9). Bottom-strand and top-strand visualisation gels were run separately. All panels for top or bottom strand were cropped from the same gel, but target-site DNA intensity was adjusted independently of products. LC indicates the loading control for product recovery with precipitation and gel loading. (d) Detection of 3’ and 5’ rDNA-insertion junctions by PCR. Detection of an rDNA-inserted transgene 3’ junction indicates successful cellular TPRT, while detection of a transgene 5’ junction indicates stable insertion. Sizes of truncated 3’UTRs used for this assay are in parenthesis. Expected product sizes are listed below each gel and those expected products are denoted with a black arrowhead.

The first functional test for our selected R2p used purified recombinant proteins for *in vitro* assays of 28S rDNA target-site nicking and TPRT. As a standard for comparison, we used the well-characterised TaGu protein with robust TPRT and site-specific gene insertion capability in human cells *(9, 30)*. As done previously, R2p was expressed from a synthetic ORF using transient transfection of human HEK293T cells and purified by FLAG antibody resin *(9)*. Also, as done previously, a template RNA was *in vitro* transcribed that harbours the species-matched R2 3’ UTR 3’ end and a 3’ tail complementary to the nicked primer (SI Table 8) *(9)*. Annealed target-site oligonucleotides had 5’-end dye labels to detect first-strand (bottom strand) nicking and cDNA synthesis as well as second-strand (top strand) nicking (Fig. 4b-c, schematics at top). We tested R2p activities both with intact target-site duplex (Fig. 4b) and also with bottom-strand “pre-nicked” target-site duplex (Fig. 4c) to evaluate TPRT with or without the requirement for first-strand nicking.

Several recombinant R2p demonstrated robust first-strand nicking and TPRT comparable to the positive-control TaGu R2p, including lizard PBla, turtle PlaMe, and avian AgTri (Fig. 4b). Fish LaMa and NePa, snake CroHe, and salamander DeFu had weak activity on the intact target-site substrate (Fig. 4b). LaMa and DeFu TPRT activity was improved using pre-nicked target site DNA, and weak TPRT by gharial GaGa and Tuatara SPun became detectable (Fig. 4c). Second-strand nicking activity tracked with first-strand nicking activity (Fig. 4b-c, lower panels). Overall, within this sampling, we conclude that most tested R2p have nicking and cDNA synthesis activities. However, weak first-strand nicking limits TPRT by LaMa, DeFu, GaGa, and SPun, evidenced by increased TPRT with pre-nicked target-site duplex.

Our second functional test used a version of the PRINT protocol for R2p-mediated transgene insertion *(9, 36)*. In brief, a proliferatively immortalised RPE-1 human primary cell line was co-transfected with mRNA encoding an R2p and a template RNA with the species-matched R2 3’ UTR 3’ end and primer-complementary 3’ tail. We detected successful genome insertion at the R2 28S rDNA target site by PCR. Again, we used TaGu as a positive control, and we compared template RNAs with native uridine or the 100% pseudouridine substitution that enhances TaGu-mediated genome insertions *(9, 36)*. Template RNA transfection without R2p mRNA was the negative control.

Across the R2p panel, successful cellular TPRT was detected by transgene 3’ junction formation when R2p mRNA and template RNA were co-transfected (Fig. 4d, top panel of each set; first sample in each panel is parental cells alone, and the next two are template RNA without R2p mRNA as negative controls). Efficiency of transgene insertion generally mirrored *in vitro* TPRT activity on intact target-site duplex (Fig. 4b). PBla, PlaMe, and AgTri were most active; NePa, DeFu, and SPun appeared not active in cells, while LaMa, CroHe, and GaGa were weakly active (Fig. 4d). Curiously, CroHe and GaGa preferred template RNA with uridine rather than pseudouridine, which we speculate could result from a modified-nucleotide influence on RNA folding or binding to R2p. We also detected rDNA-insertion 5’ junctions (Fig. 4d, bottom panel of each set) that compared to the TaGu positive control were robust for PBla and AgTri, with PlaMe, CroHe, and GaGa also showing detectable levels of stable transgene insertion compared to the negative controls (first 3 samples of each panel). We conclude that when assayed in human cells, many tested R2p retained the functionality for gene insertion that is essential to native retrotransposon mobility.

## Discussion

In this study, we curated R2 retrotransposons from 300 species across 12 key phyla. The increasing availability of genomic sequence information in long reads, short reads, and genome assemblies enabled our expanded R2 curation to be optimised to retrieve full-length R2s. In this study, we show that R2s have persisted most steadily over Metazoan evolution in two lineages, D and A (Fig. 5 summarises the overall inheritance patterns of lineages D and A across major animal lineages). The sheer diversity in terms of species where R2s are found, as well as the diversity of R2p structural variation, were underappreciated from previous studies and even by our expectations at the launch of this work.

**Figure 5:**
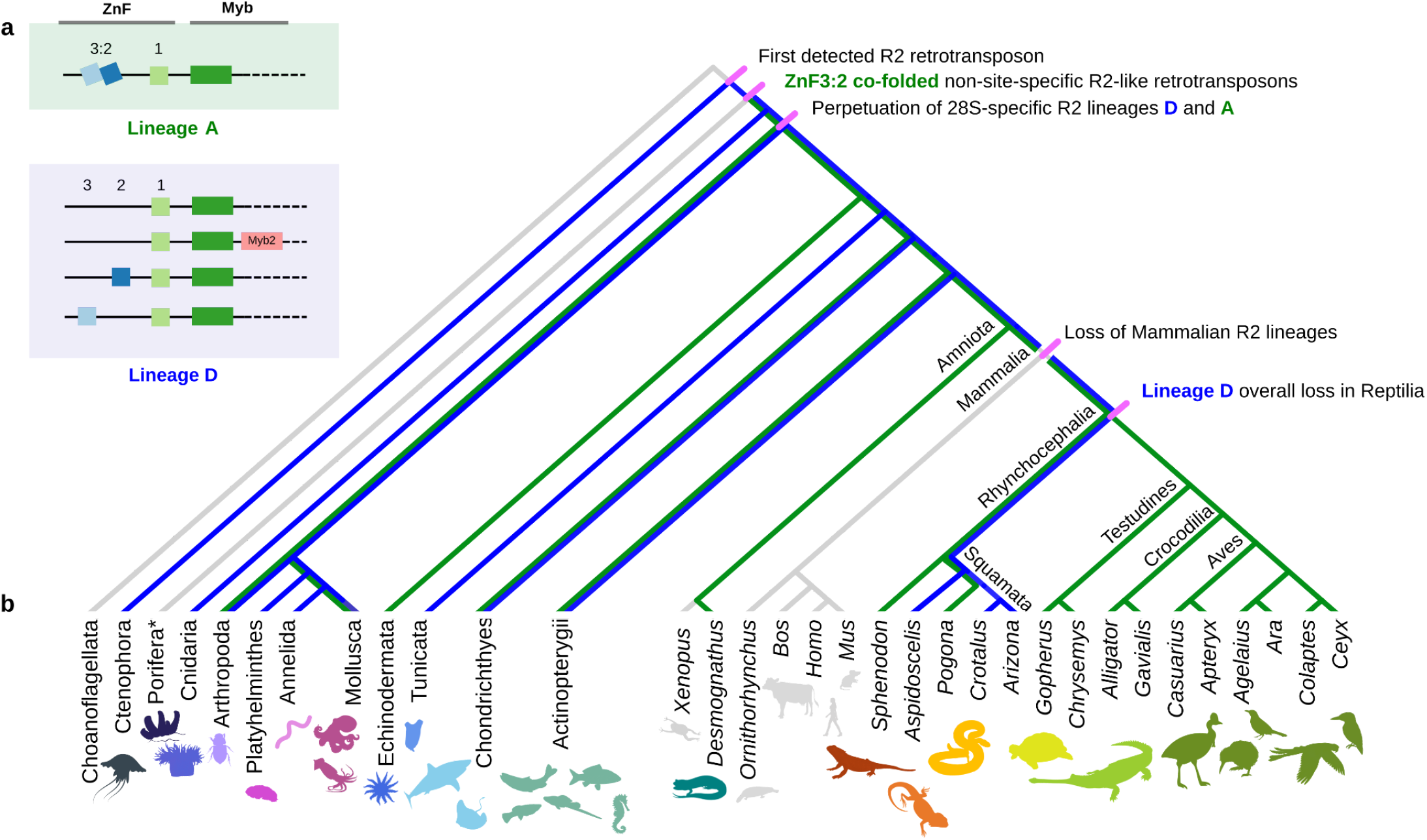
Summarising observed R2 inheritance in Metazoa. (a) Key outlining different ZnF and Myb configurations present in R2 lineages. Lineage A has co-folded ZnF3:2, while lineage D generally does not. Superimposing persistence of R2 lineages D and A in key phyla across Metazoa. Branch lengths are not on an evolutionary time scale. (*) The placement of Porifera non-site-specific R2 elements into either lineage D or A is ambiguous.

We used biochemical assays to evaluate the activities of R2p across a spectrum of phylogenetic groups and domain architectures, focusing on groups other than those previously sampled. The parallel assay conditions for each R2p do not attempt to capture physiological differences in the organisms, and therefore, negative findings do not necessarily indicate a lack of biological function. Despite this departure from the physiological context, we demonstrate that the D-lineage fish LaMa R2p with an entirely unanticipated Myb domain duplication retains function for 28S rDNA insertion. How the evolutionary innovations of N-terminal domain architectures and ZnF sequence signatures influence insertion efficiency and the fidelity of site-specific insertion remain to be investigated with more directed structure/function assays.

As would be anticipated from previous work, the host tree and R2 tree are not congruent. However, this incongruence is not predicted simply by the number of N-terminus ZnFs. Within a larger complexity, two especially successful lineages arose. We propose that the widespread survival of lineage A occurred with the preservation of 3 N-terminal ZnFs, including the co-folded ZnF3:2, a feature that may have evolved in the common ancestor prior to the split of Porifera and other animals. Co-folded ZnF3:2 may confer an advantage in enhancing as well as coordinating the recognition and positioning of template RNA and target-site DNA *(30, 37)*. We propose that widespread lineage D survival was founded on an ancestral R2p with ZnF1-Myb specific for the R2 rDNA target site, which in several sub-lineages gained an additional ZnF or Myb domain. Even considering only lineage D R2 with streamlined ZnF1-Myb, several sublineages arose that maintained independent mobility, especially notable in Cnidaria and Tunicata (see Fig. 2d). The distinct adaptations of lineage D and A RT-domain motifs (see Fig. 3e) suggest that many RT domain features co-evolved with changes in N-terminal domain architecture. One example is the change in RT motif 6a contacts to target site DNA, which are sandwiched between contacts made by ZnF1 and Myb and are different in lineage D BoMo versus lineage A TaGu and PlaMe *(30, 34, 37)*.

Tandemly repeated gene arrays, such as rDNA, undergo non-allelic homologous recombination, which can cause duplication and deletion of sequences *(38)*. Ribosomal DNA arrays are prone to copy number variation and, therefore, potential loss of rDNA-embedded R2 *(39)*. In some species, R2 persistence may be due to their ability to reverse rDNA copy number loss *(40)*. Long-term evolutionary persistence has given rise to the apparent patterns of lineage D and A inheritance (Fig. 5), with either lineage apparently lacking in some phylogenetic groups. This can be explained by the differential retention of shared ancestral lineages rather than by horizontal transfer (HT) *(41)*. While potential R2 HT has been suggested for *Bacillus* stick insects facilitated through their complex reproductive systems (*42-44)*, we did not identify other hallmarks of recent HT (e.g. unusually high sequence similarity between divergent species; SI Table 9 shows percentage identity matrix between R2 sequences). Although we do not exclude the possibility of ancient HT, our observations can be explained more simply by stochastic loss of lineages. In the future, domain architecture could be another approach to investigate ancient HT in early Metazoan evolution. For example, snake lineage D retrotransposons share the same ZnF configuration with Platyhelminthes and certain fishes and share a higher sequence identity to each other than with other reptiles (SI Table 9).

While previous studies suggest that the origin of R2 predates Metazoan evolution, the nature of the precursor non-LTR retrotransposon remains unclear. With the present R2 distribution, we hypothesise that R2 emerged from non-LTR retrotransposon(s) with some degree of target site specificity, which subsequently adapted to recognition of the 28S rDNA locus. Porifera R2-homologous retrotransposons are peculiar as they have an R2p domain architecture but have flanking sequences more like those of Utopia retrotransposons, making them appear as an ‘in-between’ state separating R2 and Utopia. The TaGu and PlaMe ZnF3:2 co-fold *(30)* can also be discovered in some Utopia elements in ZnF2:1 (SI Figure 14), however we do not know if there have been any ZnF domain swaps between Utopia and R2s. Based on this, we might conclude that the precursor to R2s was likely derived from a common ancestor shared with Utopia before it gained specificity for the 28S. However, Ctenophora R2s use the 28S rDNA target site, and the current model for Metazoan evolution places Ctenophora as the sister group to other animals rather than Porifera *(45)*. We suggest a model for R2 evolution in which rDNA site-specificity was acquired early, before divergence of Ctenophora and Porifera, and largely retained through selection for rDNA copy number maintenance. A loss or change of site-specificity would then have occurred in the Porifera branch of R2-homologous retrotransposons.

R2s in general and lineages D and A in particular have been diverging for hundreds of millions of years. Evidence for an early emergence of lineage D is in the clustering of ctenophore R2s with lineage D (Fig. 1). Also, ctenophore R2p have a general topology of ZnF1-Myb contacts with target site that is similar to BoMo (SI Fig. 1). Porifera R2-homologous proteins have weak support values in lineage D when the tree is rooted with a Group II intron protein (Fig. 1), yet are well supported within lineage A when the R2 ORF tree is rooted by ctenophore R2 (SI Fig 6.2-3). We propose that the ZnF3:2 co-folding observed in some Porifera R2-homologous proteins emerged in the common ancestor of Porifera and other animals, and that ancestral R2p gave rise to the highly successful lineage A. Lineage A is currently widespread from Arthropoda to Aves (Fig. 2c), but whether that reflects later emergence of lineage A than lineage D or lineage A loss in early-branching phylogeny cannot be resolved by our extant R2 inventory. Because purified R2p of both major lineages show the same target-site positioning of first- and second-strand nicking, a common origin of the RT-EN core seems likely. However, lineages D and A differ in ZnF1-Myb positioning on upstream DNA (SI Fig. 1), raising the possibility that lineage D ZnF1 was replaced by the gain of 3 ZnFs from an R2-homologous protein with co-folded ZnF3:2 (30, 37). The utility of the co-folded ZnF3:2 module for coordinately binding template RNA and melted duplex DNA at the nick site may have aided the success of lineage A at the expense of lineage D in turtles, crocodilians, and avians (Fig. 5) *(30, 37)*. Understanding the foundation of lineage A success is a high priority for further exploration of the features of avian R2p that confer high activity for transgene insertion into human genomes *(9)*.

## Materials and Methods

### R2 discovery pipeline

We conducted online BLASTN+ and TBLASTN searches using reference R2 query sequences (Clade A, B, C, and D R2s were used initially, and then species-specific queries were used) against the genome assembly of any species of interest (sensitive search, word size = 7) (46). If the top hit was a full-length R2 flanked by 28S rRNA, the sequence was annotated as an R2. If the top hits were truncated or appeared to be misassembled, and both long and short-read data were available, we downloaded the long and short reads from the SRA database (https://www.ncbi.nlm.nih.gov/sra). See SI Fig. 12 for full parameters and workflow. If there were no significant hits, we did not investigate that species further.

Using long-read sequences, a local sensitive BLASTN+ search was performed with reference R2 queries. We processed the short reads through FastQC, and adaptors were removed with fastp (http://www.bioinformatics.babraham.ac.uk/projects/fastqc) *(47)*. Long-read sequences containing R2s were error-corrected with short-reads using Ratatosk (default) *(48)*. We checked the quality of error correction with IGV *(48, 49)*. R2s were re-extracted from the error-corrected long reads using a local sensitive BLASTN+. Top hits were filtered for sequences exceeding 4200 base pairs with flanking 28S rRNA target sites and were annotated as full-length R2s. If the sequences were truncated, we annotated them as partial-length R2s. We translated R2 ORFs using ExPASY and NCBI ORF Finder *(50, 51)*. We used NCBI Conserved Domain Search to identify key domains *(52)*. We searched our curated R2 against the Repbase repeat masking function to confirm that we had correctly annotated our sequences *(53)*. Finally, we generated a percentage identity matrix based on nucleotide sequences to visualise R2 sequence similarities with MUSCLE *(54)*. R2 sequences from *B. mori, T. guttata, T. guttatus, T. castaneum, N. vitripennis, L. polyphemus, D. mercatorum* were used from previous studies *(19, 23, 55-57)*.

### Phylogenetic tree-building

We aligned R2 ORFs that were longer than 900 amino acids with MAFFT v7.490 (Auto model selection) and trimmed the alignment with ClipKIT *(58, 59)*. We manually checked each alignment generated with AliView *(60)*. We used IQTree v2.0.3 for tree reconstruction with 20 maximum likelihood trees and 1000 bootstraps *(35)*. ModelFinder was used to obtain the best-fit model for tree building (-m MFP) *(35, 61)*. We reduced the number of Amphibia R2s in all phylogenetic trees from 37 to 3 representative sequences. We did several iterations of tree building to test topology based on domain partitions (Partitioned analysis for multi-gene alignments) with MFP+MERGE: 1) complete R2 ORF, 2) RT, 3) ZnF-Myb, 4) RLE, and 5) Thumb-ZnF *(35, 62)*. To check our tree topology for reliability, we used RaxML *(63)* (SI Fig. 4). We visualised and edited our tree using iTOL: Interactive Tree of Life and TVBOT *(64-66)*.

We used TimeTree to build a species tree representing the major classes of organisms with R2s *(54, 66)*. This included the following: Amphibia, Arthropoda, Cnidaria, Ctenophora, Echinodermata, Actinopterygii, Chondrichthyes, Mollusca, Platyhelminthes, Porifera, Reptilia (Aves, Rhynchocephalia, Squamata, Testudine) and Tunicates.

### Protein structure prediction and sequence comparison

For protein structure prediction, we used the AlphaFold3 web server (https://alphafoldserver.com/) *(67, 68)*. Predictions were done for both protein alone and protein plus target-site DNA oligonucleotides. We used ChimeraX *(69)* v1.8 for structural analysis and alignments, including the protein fold visualisations shown. To search for structural similarity to the lineage A co-folded ZnF3:2, we used PlaMe ZnF3:2 as a query to perform a Structure Similarity Search following the recommended pipelines by RCSB.org (https://www.rcsb.org/) to query for both structures within the Protein Data Bank (PDB) and those available in other public data resources (e.g. AlphaFold or RoseTTAFold predictions). We manually examined polymer entities that ‘matched’ the query, which in all cases lacked the R2p ZnF3:2 co-folding. Cofolded ZnFs that structurally matched lineage A cryo-EM structures were only detected when we individually predicted the protein structures of apparent full-length Utopia elements described previously (SI Fig. 14) (*27)*.

We used the BLAST+ position-specific scoring matrix to calculate relative similarity *(70)*. Each predicted R2p RT was incorporated into a PSI-BLAST database and queried using BoMo and TaGu RT sequences previously validated to be active R2p *(9, 12)*. The alignment bitscores were used to visualise the similarity of each R2 to the lineage D and A representatives in Figure 3a-d (also see SI Fig. 9). To infer site-specific rate of evolution for R2p amino acids (Fig. 3e), we used IQTree (LG+R6, --rate) *(35)* (IQTree; http://www.iqtree.org/doc/Advanced-Tutorial). We extracted the numerical value for the conservation of each residue and mapped it against the RT sequence. We then annotated the position of RT motifs (0 to 7) *(33)*.

### Cell culture

RPE-1 hTERT cells were grown in DMEM/F12 (Gibco), and HEK293T cells were grown in DMEM (Gibco). All media was supplemented with 10% fetal bovine serum (FBS; Seradigm) and 100 μg/mL Primocin (InvivoGen). Cells were cultured at 37 °C under 5% CO2. All cells were tested for mycoplasma contamination, and human cell lines were validated by short tandem repeat profiling (Promega, catalogue no. B9510).

### Protein expression, purification, and detection for biochemical assays

HEK293T cells were transiently transfected with pcDNA3.1 plasmids with synthetic ORFs encoding proteins N-terminally tagged with a single FLAG peptide (SI Table 8) used for affinity purification as previously described *(9)*. In brief, cell pellets were resuspended in HLB (20 mM HEPES pH 8.0, 2 mM MgCl2, 200 μM EGTA, 10 % glycerol, 1 mM DTT, 0.2 % protease inhibitor cocktail (Sigma, catalogue no. P8340), 1 mM PMSF). After 5 min on ice, cells were lysed by three cycles of snap freezing in liquid nitrogen and thawing in a room-temperature water bath. Lysate was brought to 400 mM NaCl, gently vortexed, and placed on ice for an additional 5 min; then, NaCl was lowered to 200 mM and 0.2 % NP-40 was added prior to centrifugation. Clarified extract was incubated with FLAG resin (Sigma, catalogue no. A2220) at 4 °C for 2 h. Resin was washed with the same buffer and protein eluted with 50 ng/μL 3xFLAG peptide (Sigma, catalogue no. F4799) at room temperature for 1 h. The slurry was aliquoted, snap-frozen, and stored at -80 °C. Immunoblots used anti-FLAG antibody (Sigma, catalogue no. F1804, 1:3000) and Alexa Fluor 680 goat anti-mouse secondary (Thermo Fisher, catalogue no. A21057, 1:2,000) detected by LI-COR Odyssey.

### RNA synthesis

R2 mRNAs and template RNAs were synthesised using T7 RNA polymerase as previously described *(9, 36)*. In brief, mRNAs were made using plasmid template linearised with BbsI-HF (NEB) for 4 h at 37 °C, purified, and transcribed with AG Clean Cap (TriLink, catalogue no. N-7113) per the manufacturer’s protocol, with complete substitution of uridine with N1-methylpseudouridine (TriLink, catalogue no. NC1443155). The mRNA UTR sequences are from the BioNTech COVID-19 vaccine mRNA as previously in this work, followed by a plasmid-encoded 30-nucleotide poly-adenosine tail *(9)*. Transcription reactions were incubated at 37 °C for 2 hrs, followed by the addition of 2 μL RNase-free DNase I (Thermo Fisher, catalogue no. FEREN0521). Product RNA was purified by desalting with an Illustra Probe-Quant G-50 Micro Column (Cytiva, catalogue no. 28903408) followed by phenol-chloroform-isoamyl alcohol (PCI; Thermo Fisher, catalogue no. BP1752I-100) purification and precipitation with a final concentration of 2.5 M LiCl. After washing with 70 % ethanol 2-3 times, RNAs were resuspended in 1 mM sodium citrate (pH 6.5). Truncated R2 3′ UTRs were transcribed from PCR-amplified DNA template using uridine (NEB) or pseudouridine (TriLink, catalogue no.) with the HiScribe T7 Kit (NEB, catalogue no. E2040S) according to the manufacturer’s instructions for RNAs <300 nt. Purification was the same as for the mRNA with the following exceptions: following RNA synthesis 1 μL DNase RQ1 (Promega, catalogue no. M610A) was added to remove PCR template; following the RNA PCI purification step precipitation was done with a final concentration of 0.3 M sodium acetate (pH 5.2) and 3 volumes of 100 % ethanol; and RNAs were resuspended in water. RNA concentrations were determined by NanoDrop and integrity verified by denaturing urea-PAGE with direct staining using SYBR Gold (Thermo Fisher, catalogue no. S11494). Sequences for mRNA and template RNA transcription are provided in SI Table 8.

### TPRT assays with purified protein

IR700 labelled target site oligos were ordered from IDT (SI Table 8). Complementary strands with only one of the two strands labelled were annealed by heating to 95 °C and cooling by 1 °C per min. TPRT reactions were assembled in 20 μL with final concentrations of 25 mM Tris-HCl pH 7.5, 75 mM KCl, 5 mM MgCl2, 10 mM DTT, 2 % PEG-6K, 5 nM target-site duplex, 0.6 μM template RNA, 50 μM dNTPs and approximately 3-10 nM R2p then incubated at 37 °C for 30 min before heat inactivation at 70 °C for 5 min. RNaseA (Thermo Fisher, catalogue no. FEREN0531) was added to a final concentration of 0.5 mg/mL and incubated at 55 °C for 30 min before dilution with 80 μL of stop solution (50 mM Tris-HCl pH 7.5, 20 mM EDTA, 0.3 % SDS) spiked with a 36-nucleotide synthetic loading control (LC) oligonucleotide. Nucleic acid was purified by PCI extraction and ethanol precipitation, resuspended in 5 μL of water, and supplemented with 5 μL of formamide loading dye (95 % deionised formamide, 0.025 % w/v bromophenol blue, 5 mM EDTA pH 8.0). The sample was heated to 95 °C for 3 min and then placed on ice before loading a 9 % urea-PAGE gel. After electrophoresis, the gel was imaged by Typhoon Trio (Cytiva). The gel was then stained with SYBR Gold and imaged again to visualise the ladder and LC.

### Transgene insertion in cells

PRINT was performed largely as previously described *(36)*. RPE-1 hTERT cells at 50% confluency were replated at 400,000 cells per well in twelve-well plates. Cells were reverse-transfected with mRNA and template 3′ UTR RNA using Lipofectamine MessengerMAX at ½ mass/volume ratio as per the manufacturer’s instructions. 0.5 μg total RNA mixture was transfected per well of a twelve-well plate, and mRNA:template molar ratio was 1:20. Cells were collected 20-24 h after transfection and frozen as cell pellets. Genomic DNA was isolated by treatment with RNase A and Proteinase K, extraction with PCI, and precipitation as previously described *(9)*. For PCR, 100 ng gDNA was used in a 25 μL reaction with Q5 DNA polymerase (NEB). PCR primer sequences are listed in SI Table 8. PCR was as follows: 98 °C, 3 min, (98 °C 10 sec, 65 °C, 30 sec, 72 °C, 30 sec/1 kb) 5 times with annealing temperature decreasing by 1 °C per cycle, (98 °C 10 sec, 60 °C, 30 sec, 72 °C, 30 sec/1 kb) 25 times; 72 °C, 2 min. PCR products were analysed on 1% agarose gels containing GelRed (Fisher Scientific, catalogue no. NC9594719) and imaged using the Bio-Rad gel doc XR+-imaging system.

## Acknowledgments

We thank the Adelson and Collins lab for their support for this collaborative project. We especially thank Terry Bertozzi, Alex Stuart, Zhenglong Du, and Rick Tearle for their input when preparing this manuscript.

## Author contributions

Conceptualisation: N.T.H, K.C., A.E.S, D.L.A,; Methodology: N.T.H, B.V.T, A.R.V.; Investigation: N.T.H, B.V.T, A.R.V.; Visualisation: N.T.H, B.V.T, A.R.V.; Supervision: K.C, A.E.S, and D.L.A.; Writing–original draft: N.T.H, K.C, D.L.A.; Writing–review: all authors.

