## Supplementary FIgures for "Lineage-Specific Evolution, Structural Diversity, and Activity of R2 Retrotransposons in Animals"

#### Supplementary Figure 1

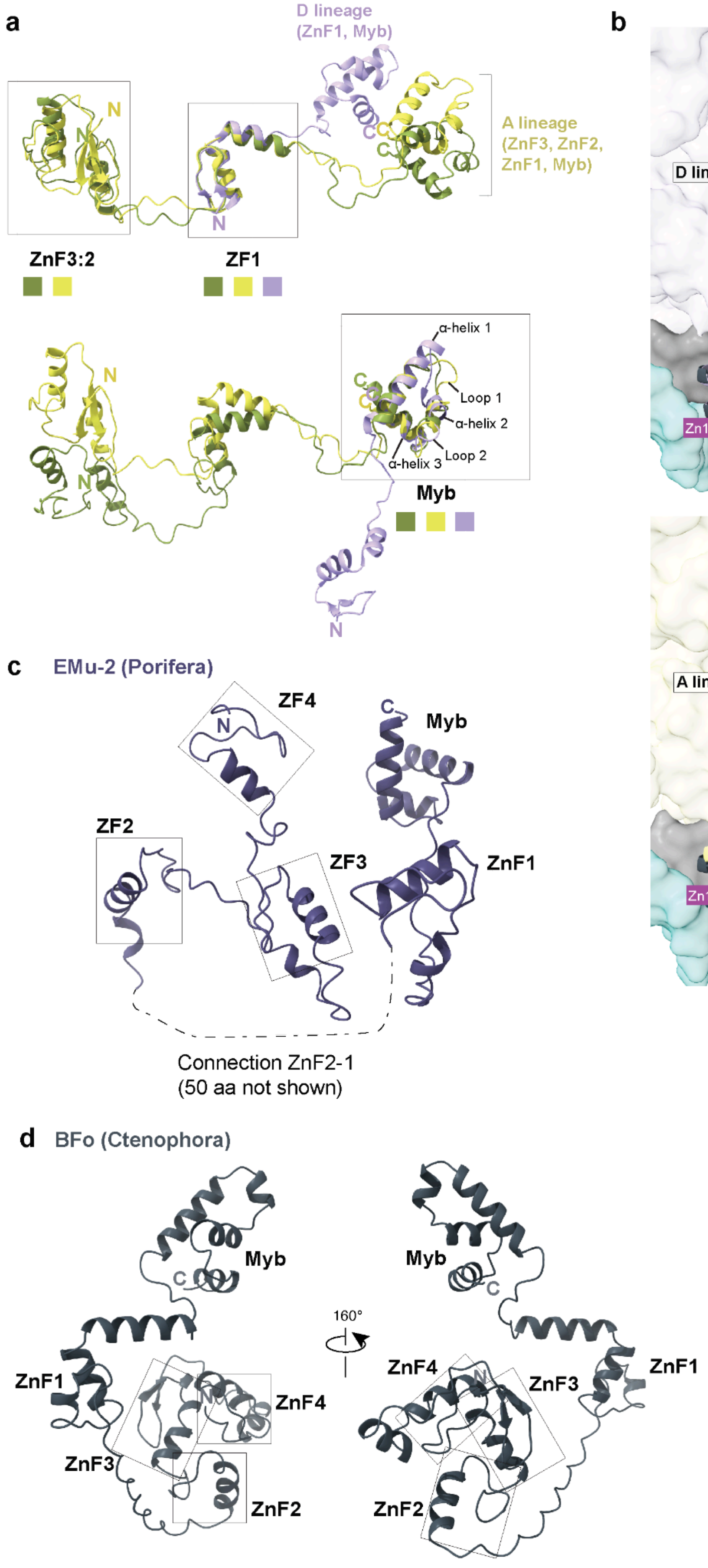


**Figure S1:** Structural arrangements of R2 N-termini to classify folds of novel protein conformations. (a) R2 structural references of the N-termini studied from R2 cryo-EM structures of complexes bound to the same DNA substrate, including A lineage (R2Tg - TaGu, R2Pm - PlaMe) by our group [1] and D lineage (R2Bm - BoMo) by others [2]. Three R2 N-termini are superimposed on each other and each box highlights the structural regions that best align. Coloured squares below each domain represent the matching colours for each R2 reference (TaGu is green, PlaMe is yellow, and BoMo is purple). Top panel: ZnFs across elements can be aligned, with the exception of BoMo which only has ZnF1. Bottom panel: Myb domain requires a local alignment to confirm similar fold architecture containing three expected alpha helices connected by two loops. (b) Conformation of Ctenophore ZnF1-Myb bound to DNA relative to R2 lineages using BoMo (D lineage, top panel) and PlaMe (A lineage, bottom panel) as structural references. All ctenophores examined (n=4) with AlphaFold (AF) predictions, BMi as the representative shown, have a D-lineage-like Myb conformation, where Myb is spatially closer to the RT domain. (c) A structural representative (AF structure) of independently folded ZnFs where ZnF2-ZnF4 (boxed) do not co-fold. (d) Two orientations of an AF structure for BFo showing a distinct fold arrangement where ZnF2-ZnF4 (boxed) have shorter connections between them than model R2s in (a).

#### Supplementary Figure 2

**>WFMF01000099.1:20329-29541 Beroe forskalii isolate Bf201606 sca99**

AATAGGACTTTGGTCTTATTTTGTTGGTTTCCGAGACCGAAGTAATGATTAATAGGGACAGTTGGGGGCATTCGTATTTCATTGTCAGAGGTGAAATTCTTGGATTTATGAAAGACGAACTTCTGCGAAAGCATTT
GCCAAGGATGTTTTCATTAATCAAGAACGAAAGTTGGAGGCTCGAAGACGATCAGATACCGTCCTAGTTCCAACCATAAACGATGCCGTCTGCGGATCGGAGGTCGCTAACGTAAGGCTCCTTCGGCACGCTATGA
GAAATCAAAGACTTCGGGTTCCGGGGGGAGTATGTTCGCAAGAATGAAACTTAAAGGAATTGACGGAAGGGCACCACCAGGAGTGGAACCTGCGGTTTAATTTGACTCAACACGGGAAAACTCACCAGGTCCAGAC
ATAGGAAGGATTGACAGATTGAGAGCTCTTTCTTGATTCTATGGGTGGTGGTGCATGGCCGTTCTTAGTTGGTGGAGTGATTTGTCTGGTTAATTCCGTTAACGAACGAGACCTTAACCTGCTAAATAGTGACACG
GTTATTTTAACCGTGGTTCACTTCTTAGAGGGACTATCGCTTTCAAAGCGATGGAAGTTTGAGGCAATAACAGGTCTGTGATGCCCTTAGATGTTCTGGGCCACACGCGCGTTACACTGATGAAGCCAGCGAGTAT
ATCGCCTTCACCGAAAGGTGCGGGTAATCTTGTGAAACTTTATCGTGCTGGGGATAGACCCTTGCAATTATTGGTCTTACTAATCGAACCATCTAGTAGCTGGTTCCCTCCGAAGTTTCCCTCAGGATAGCAGGAG
CTCATTCCCAGTTTTATCAGGTAAAGCGAATGATTAGAGGCCTTAGGGATGAAACATCCTTAACCTATTCTCAAA**CTTTAAATGGGTAAGATGTC**CGACTTGCTTAATTGAAGCCGGGCC

R9 target site

ACGAATGGGAGCTCCTAGTGGGCCATTTTTGGTAAGCAGAACTGGCGATGCGGGATGAACCGAACGTTTGGTTAAGGCGCCTAAATCGACGCTCATCAGATCCCACAAAAGGTGTTGGTTGATATAGACAGCAGGA
CGGTGGCCATGGAAGTCGGAATCCGCTAAGGAGTGTGTAACAACTCACCTGCCGAATCAACCAGCCCTGAAAATGGATGGCGCTCAAGCGTCGTGCCTATACCAGACCGTTTGAGCAAATGCTAAGCTCGGACGAG
TAGGAGGGCGTGGGGGTCGTGAAGAAGCCTCTGGCGTGAGCCTGGGTCAAACGGCCTCTAGTGCAGATCTTGGTGGTAGTAGCAAATATTCAAATGAGAACTTTGAAGGCCGACGTGGAGAAGGGTTCCATGTGAA
CAGCAGTTGGACGTGGGTTAGTCGATCCTAAGAGATAGGGAAACTCCGTGTTAAAGTGCCCGATCTACGACTCACGTCAATCCGGGCCGCCTATCGAAAGGGAATCGGGTTAATATTCCCGAACCGGAATGGGGGG
ATTCTGTCTTTCGGGACAGAAATGCGGTAACGCAAACGAACTGGGAGACGTCAGCCGGGAGCGGAAGCAGAGTTCTCTTTTCTGTTTAACAGACAGTCACCATGGAATGGGATTATCCCGAGATATGGTTCAGTGT
CTGGTAAAGCACCGCGCTTCTTGTGGTGTTTCCTGCCTTCCGGATGGCCCTTGAAAATCCCAGGGAGAGAATAAACTTCTCATCCGGTCGTACCCATAACCGCAGCAGGTCTCCAAGGTTAACAGCCTCTGGTCGA
TAGAGCAATGTAGGTAAGGGAAGTCGGCAAAATAGATCCGTAACTTCGGGAAAAGGATTGGCTCTAAGGGTTGGGTCTGTCGGGCTGCAGCCCGAGACCTCGAAGCCTTGGGTGGACTGTCTAGAGTAAGGTTCGC
CGAGCTCCGGATGGGCCGTCCTCGGCATCGAGGAGGAAGGCTGCAGCTTCGTCTCCGGACATTTCGGCAGACGACTAACAACCAACTTAGAACTGGTACGGACAAGGGGAATCCGACTGTTTAATTAAAACAAAGC
ATTGCGATGGCCGGAAACGGTGTTGACGCAATGTGATTTCTGCCCAGTGCTCTGAATGTCAAAGTGAAGAAATTCAACCAAGCGCGGGT*CCTTAAAATTGGAATCGAAGAAGTGGGACATATGAGAAGGGAGAAT

Shift in insertion site

GAACGCTGACTTAGGCAAGAAAGTTGAGATGGAACGAAAGTAAAGAGTGTGACCCTCTTCTTCCCTCCAGCGAGTCACTGATACTTGTTCTTTAAGCCCTTGACGAAGTAAGATGAATACCTTTATACGTTATTGT
CATTTAGATGCGAGACAAGGCTGCGAGCAAGACAAGGCAAGCAATAGACCGTGGGTTCGTACCCAGTGACACCAGGGAAAAGAAAAAAAAAAAATGTCAATCTCGTGTAAGGTTATTTATTTTGGTACGGGCGGAG
AGATAGACGGCCAGGTCTACCCGTACAGGTCCGGCCAACGACTTGTCGTTCAACGGATTGCTTAGTCTAAGGAAGTCCCAATCCTGTCCTACAAGCGGAACCGCTGTTAGAGTGCGACATGGGGCACATGTGCAAC
CTCATGACAATCTCCGTCCTGAGATGGCTTCTCGGGTATGCGAAAACAGAGTGGAAGGGGCTAGTGAGCCTTGCTGGGCTCATCTCTTATTTTTAACAAG**ATG**GAAGAAAAAAATAATGAGACTGCCGGCCGAACG
GCTAGCAGTGATAAAACCAGCAAGAGCTCGGCTTCAGGTTCAGCTGCACGCGCCGCTGTGACTCCTGCACCCAGTAGAGCTTGCCTCTTCTGTGGTAAGGCCTTTAAACTCAGAGGCCTGGCGGGACACACGCGTA
GATGCGAGTCGAATCCTGAATGGTTGAACGAGAAAATTCATAGCTCCAGGTGGGACTTCAGTGGTAGATCGCCCGTAAACTGCGCTCGTTGTGATCTTCATATGAAAGCCACATGTTATATCGGGCACTTTCCGGA
TTGCATCGCGACTAACGGATGCTTCCGATGTGGAAATCAAGTGCCCGTGGCTGATTATGTTGAGCACTTTCAAGTTTGTGTGAGGACGATTGATCCCGCTGAACCACCAACCCCACAGGTGGTTATACCTGAGGTA
CAGAAGGTGACGTGCCCTTATTGCCCCGCTGCTCTTCAAGCTAAATTTAGCTTTAAACCTGGGCGAGGCCTTTCAATGCATGTCAAGAAGAAGCACCCTAAAGAGTGGAATGATTACCAACTGAACAGGATGGAAC
TTTACCCCTCCCAGTACATCTGGCGGGAGGGTGACGATGAACTGATTTGCCAGGCGATCGAGGAGTATCAGTCCTCAGCTCCCGCGGTTAAAAAGGCGTTTGGCATCAATCAGTTTATCCAATTCAGATCCTTCCC
ACATTGCACTCTTGACAGCATAACGTGTCACAGGAAAACTTCTTCTTTTAAAGAATATGCGAAAGACCGAGCCAAACGCATAGCTGATCAACAGGCTGTGGCTGAGTCACGAGCGGAAGAGTCACTTCCCAACGAG
GATGGAGGCCACTTCGAAGAGGTAGAGATACTCTCTGAGAAGGAAAAATCGGCCATCGCTGCCAAACCTTTTGGCAGTGAAATTCTTGAGATCCATCAGCATCTTGTGAACAGGAGATGGGATACAGCATGTAGTG
CAGCTAAGTTAGTGTTTGAAGAGCTTACTTGTAAGTTCGGAAAGCCTAATATCCCGGTTCATAAAGTGAAACAAGGTGCGGATCCCAGGCCTCCAGGGAAAAAACCCACACGCAAGTCTAAAAAGAGGGCAAATAG
GCGACGACTCAAGCCCAGCAAGGAAAAGAGAAGGGAAAGTTATGCAAAGGTACAGAAGAGGTGGAGCAAGAAAAGGGTGGACGTCATCAAATCTATCTTGGACGGCAAATTGGAGCAAGTACAAGGAGAAAAACCA
ACGATGGAGAAGCAGCACGATTACTGGAAGAGTCTGTTTGAAAGGCCTAGTCCTGAAGATCCAGGTAAAGTGCCGGGGGAAAGTTCAGTACAATATGAATTGAGTGCTCCGCTGACGAAGGAAGAGGTAGCCAACG
GCTTAGGAACAGCCAAAGAGGCCTCGACCGGACCAGATGGGGTTCCTCTCTCTGCGTTAAAGGAACTTGGTTCCTTGAGTCTGTGGGTCCTGTACTGTGCGCTGTGGATGAAGTCGGAAACTCCGAAGGGTTGGAG
GGTTAGCCGGACGGTCTTAATAGACAAGACGGCGGACGAATGTGGGAAAAACCCGGAGGACTTCCGACCGATATCCATCAGTTCGTACTTTTACAGGATCTACGCGAGATCGATCAGCTCGCGCTTGACGCGAGCA
GTTCCAATCTCGAAAAGACAGCGTGGATTCATAAGGCAAGATGGTGTCAGAGACAATATCGAATTGTTCGATAATTTGGTCAAGGACGCCAAGCGAACCTTAACCCCCTTGAGCGTGGCCTTTATGGATATAAGGA
AGGCGTTTGACAGTGTTGGCCACGCGAGTATTCAGCGTGCCCTGGAATGGGCTGGTGTCCCTGGTCGGGTGAGGAGGGTCATCGAGGAACTATACACGGACTGCTATACGGAGGTTGGAGGGGGCAGGATTCGTGT
GAACAGAGGGGTGAAGCAGGGTGACCCGCTGAGCTCCTTCTTGTTCAATATAGTTCTCGAGATGGCACTCAGTAGAGTTCCACCAGGCCTGGGCATCAACTATCTGGGGCATCAACTGTTTTACATGGCCTTCGCT
GATGATATGATTCTTCTAGCTAGGAATGCCCACGTGCTACAAAGAATAGTGGACATGGTGTCAAGCGAATTAGGGTTAGCTGGTTTGGAATTTAACCACGCTAAGTGTAAAGTTCTCTCCCTGGTTACGGACCCCA
GGAATAAAGCGAGTATGATAGACACTGACGTCGAGATAGCGGTAAACGGTGATCGTATCCCACCCCTCGGGGTGTTGGATACCTATCGTTATCTAGGGATAGACGTCGGAGTGAAGGGAGTTCCCAAATTCGAACC
GCAGAAGGAACTCAAAGACTTGCTTCAGAGACTCACGAAAGCACCGTTGAAACCACATCAGCGGCTCTATGCCCTGAGAGTTCACACCATGCCTCGATTCCAGCACAAGCTTATCTTTTCGAAGGTAACTAGGAAC
GCTCTAAGCCAAATGGACTCATTAGTCAGGAAGCACGTAAGAGAGTGGCTCAAATTACCCGACGATGTCACCAAGGCGGCGCTGTACGCCGACGTTGGGTCTGGAGGATTGGGCCTCATCTGCCTCGAACGAAGGG
TACCCCTCCTGAAATATTCGAGGATAGAAAGATTACGCGATTCTGATGACGAACTAGTCCGATTGTTAATCGACCAAGAACAGGTGAACAAGCGAGTGATGCAATGGAAAGATCGTTGTAAGATGAGTGGCAAGAC
CTATCGTAGTAAAGAGGAGCTCAGGGATCTGTATAGAACCCAATTGAACTCCACAGTCGATGGGAAGGGTATTAGGGGAGATACCCGGCTAGGCAGACTATCGATGCGGTCGCTCGTTGGACCCTGCATCCCACTC
ACTCCAACTAGATACATCCATGCTATAAACGTTCGTCTTGGAACCCTACAGACTTCCGAGAGAAAGGCCAGAGGAAGGGTCTGCAACGAGAACGGCGTCCTATGTGACAAAGAGAGGGCATGTCACGCTAGGAAAG
CTGTTGCAACTCTAGGTCATATATCTCAAGTATGCCCTGTAGTACACGGTCTCAGAGTGAGAAGGCACGACCGAGTAAGAGACCGTGTTGCCAACCAACTGCAGAGCAGAAAAAAGTCTGTTGAGAAAGTACTGGT
TGAGAAGACGATAAAACTCCAGGATGGTACTGTGCTGAGACCGGACATCATTGTTCTCACTGACACTTCTATTGAGGTTATTGATGTCCAAATAAAGGCTGATATGGGAATCCGGAGGGACCTCGAGACCGACCAG
ACCGCAAAGAGAGCGAAGTACGACAGGGGCGACGTACAAACATCGATAGAACGTGAATTGTCGGTGGTTGGGATGGGGAGGCTGTATACAGTTACGGCCTTGACAATCACGTTCAGAGGGCAGCTGCCAAGACACA
CAGTTGATCTGGCTACGCGTCTTAAATTCAAGACGCTATTGCCAGAACTAGTGGCTGACGTCTTGGCAGACACGGGTAGTATGTTTGTGGTCTGGCACAGGACGACGGGAAACGCAGGAAAACCGTAGATGCCTTT
GTCT**TAG**GGATAGAAAAAGTCCCCCCTCTGCTACATCATCAGCGAGGCCAGGGACGCCAATAGTCACAGTAGGCTGGTTGTTTTACCATCAACGTAAAAGAGAACGTGTCAGTATCTTCCCACCTACTGCATAAAT
GGAACTCGATTCGACCAGGATGAGTTTGACTTGCACTACGAACGCAGGGTCACAGCTTAGTAGTGAGTTTGAACGCGATGATGACGGTGCTATGCACTGCTGATGATCTGAAGTCGCTGCTATTGCTCGAAGCAAA
CGCAGGCCACGAACATCACAGATCGTCGTATAAATGTCGTCCAGCGCAGCTTGCCCTGAAAGCGAGCTGCGGCGTCATCTGAGGAAGCCATGGCGACGGGGATGAAGCGGATGAACCATAAACATCGTCTTCGGAC
ACAAAAAAAGTAGAAAAATGCTGATCAGTACGTTCGTGGCTGATACGGCTTCCTGCGGATCCGTCACAGCCATGGTGGAATGGATGACACTGCCCTAGAAAAATCAGGGGCTCACCGCTCAGTACGGCGAGTCTTC
CACAGAGGGTGAAGCCTCAGAAGGGAGTCTCGACTCGTCCTCTCAATCCATTGGGCGAGATAGGAGTTGGCCAAACCTGACGAGAGCAGGCCGATTCCCGAGGGTACGAGGCCTCCCCCCGGGGAGGTTCTCAAAA
ATCGACGGTGAGTGACGACGCGATGAGCGTACGGACGCTATCAGTGGCCGAAAGTAGAAACTCTTCCCGATATTTTCCGTAGGTCGCAAGGCCGGTATCGGGGGAAAGCATGAATTCTTACGATTTCACAGCGAAA
GAGT***CTCTCTTAAGG*TAGCCAAATGCCTCGTCATCTAA**TTAGTGACGCGCATGAATGGATTAACGAGATTCCCACTGTCCCT

*B. forskalii* shifted and normal R2 insertion site

ATCTACTATCTAGCGAAACCACAGCCAAGGGAACGGGCTTGGCATAATCAGCGGGGAAAGAAGACCCTGTTGAGCTTGACTCTAGTCTGACTTTGTGAAAAGACATGAAGGGTGTAGCATAAGTGGGAGCGTAAGC
GACATTGAAATACCACTACTTTCATCGTTTTTTTACTTATTCGGTGAAGCGGAAGCGAGGTGCAAGCCTCACCTTCTAGATTTAAGCCGCACCTTTCGGAGCGGTGATCCGAGCCGGAGACACAGTCAGGTGGGGA
GTTTGGCTGGGGCGGCACATCTGTCAAACGATAACGCAGGTGTCCTAAGGTGAGCTCAAAGAGAACGGAAATCTCTTGTAGAACAAAAGGGTAAAAGCTCACTTGATTTTGATTTTCAGTATGAATACAAACTGCG
AAAGCATG

**Figure S2:** Nucleotide sequence of *Beroe forskalii* R2 annotated to show likely start codon (red), coding sequence (green) and stop codon (red). R9 and R2 target sites are in bold and underlined. The normal and shifted insertion sites are denoted by asterisks (black and red, respectively). 5' rDNA is in blue.

#### Supplementary Figure 3

**>BFo_R2_shifted_from_R2_targetsite**

```
MEEKNNETAGRTASSDKTSKSSASGSAARAAVTPAPSRACLFCGKAFKLRGLAGHTRRCESNPEWLNEKIHSSRWDFSGRSPVNCARCDLHMKATCYIGHFPDCIATNGCFRCGNQV
PVADYVEHFQVCVRTIDPAEPPTPQVVIPEVQKVTCPYCPAALQAKFSFKPGRGLSMHVKKKHPKEWNDYQLNRMELYPSQYIWREGDDELICQAIEEYQSSAPAVKKAFGINQFIQ
FRSFPHCTLDSITCHRKTSSFKEYAKDRAKRIADQQAVAESRAEESLPNEDGGHFEEVEILSEKEKSAIAAKPFGSEILEIHQHLVNRRWDTACSAAKLVFEELTCKFGKPNIPVHK
VKQGADPRPPGKKPTRKSKKRANRRRLKPSKEKRRESYAKVQKRWSKKRVDVIKSILDGKLEQVQGEKPTMEKQHDYWKSLFERPSPEDPGKVPGESSVQYELSAPLTKEEVANGLG
TAKEASTGPDGVPLSALKELGSLSLWVLYCALWMKSETPKGWRVSRTVLIDKTADECGKNPEDFRPISISSYFYRIYARSISSRLTRAVPISKRQRGFIRQDGVRDNIELFDNLVKD
AKRTLTPLSVAFMDIRKAFDSVGHASIQRALEWAGVPGRVRRVIEELYTDCYTEVGGGRIRVNRGVKQGDPLSSFLFNIVLEMALSRVPPGLGINYLGHQLFYMAFADDMILLARNA
HVLQRIVDMVSSELGLAGLEFNHAKCKVLSLVTDPRNKASMIDTDVEIAVNGDRIPPLGVLDTYRYLGIDVGVKGVPKFEPQKELKDLLQRLTKAPLKPHQRLYALRVHTMPRFQHK
LIFSKVTRNALSQMDSLVRKHVREWLKLPDDVTKAALYADVGSGGLGLICLERRVPLLKYSRIERLRDSDDELVRLLIDQEQVNKRVMQWKDRCKMSGKTYRSKEELRDLYRTQLNS
TVDGKGIRGDTRLGRLSMRSLVGPCIPLTPTRYIHAINVRLGTLQTSERKARGRVCNENGVLCDKERACHARKAVATLGHISQVCPVVHGLRVRRHDRVRDRVANQLQSRKKSVEKV
LVEKTIKLQDGTVLRPDIIVLTDTSIEVIDVQIKADMGIRRDLETDQTAKRAKYDRGDVQTSIERELSVVGMGRLYTVTALTITFRGQLPRHTVDLATRLKFKTLLPELVADVLADT
GSMFVVWHRTTGNAGKP
```

```
BFo        --------MEEKNNETAGRTASSDKTSKSSASGSAARAAVTPAPSRACLFCGKAFKLRGL
BMi        ------------------------------------------------------------
MLe        KLHRCCPLFQEKDSGGLNRSAS------------------------------------GV
PBa        TL----PFLKLNMNRLNNEKNS------------------------------------GA


BFo        AGHTRRCESNPEWLNEKIHSSRWDFSGRSPVNCARCDLHMKATCYIG-HFPDCIATNGCF
BMi        ------------------KNTMFNITPR---PEDSQVDRVTIDAESGLPNHAGANLLQCE
MLe        MSNTSHSKLNLK-MDNKLKTSLETPSGV---RADSIITRVRTSSNRG-EHSNGVTYPRCE
PBa        VRSTEFSV-----ADNRPSQSLRTTESH-----------------------------RCP
                                :                                     *

BFo        RCGNQ-VPVADYVEHFQVC----------VRTIDPAEPPTPQVVIPEVQKV-TCPYCPAA
BMi        WCDRLCKNKAGLTLHKRACKNNPAVGSSAGNTNDNRRINTP----PTMRSLFNCEYC---
MLe        ------QGVAPLDTHGGICDAPPQVTVPATETDKQKK----------------CEYC---
PBa        NCRKLCRSGNGLALHMKHC-------AKCYQNGDNRQEPVK----PRM----ECSIC---
                         *   *           .. .  .                *  *

BFo        LQAKFSFKPGRGLSMHVKKKHPKEWNDYQLNRMELY-PSQYIWREGDDELICQAIEEYQS
BMi        ---NTGYGTDRGLSAHISKKHIPEWNMIKMERFKADGPRQHVWREGDWEILCQGEEEHDR
MLe        ---EFTYLKPRQIGTHMRKRHPQEWNDIKRTKFLSE-KRQKRWLDEDFELLCIGQEEYLV
PBa        ---GLFFSGQRGVAIHKRKKHPAEWNETKRVEDTLS-RKKIRWTSGDRELLHLGMIEWEA
                 :   * :. *  *:*  ***  :  .       :  * . * *::  .  *
```

**Figure S3:** (Top) Translated R2 ORF from *B. forskalii*. (Bottom) MSA of all Ctenophora R2 N-terminal ZnFs. CXXC residues in the ZnFs are highlighted in green.

#### Supplementary Figure 4

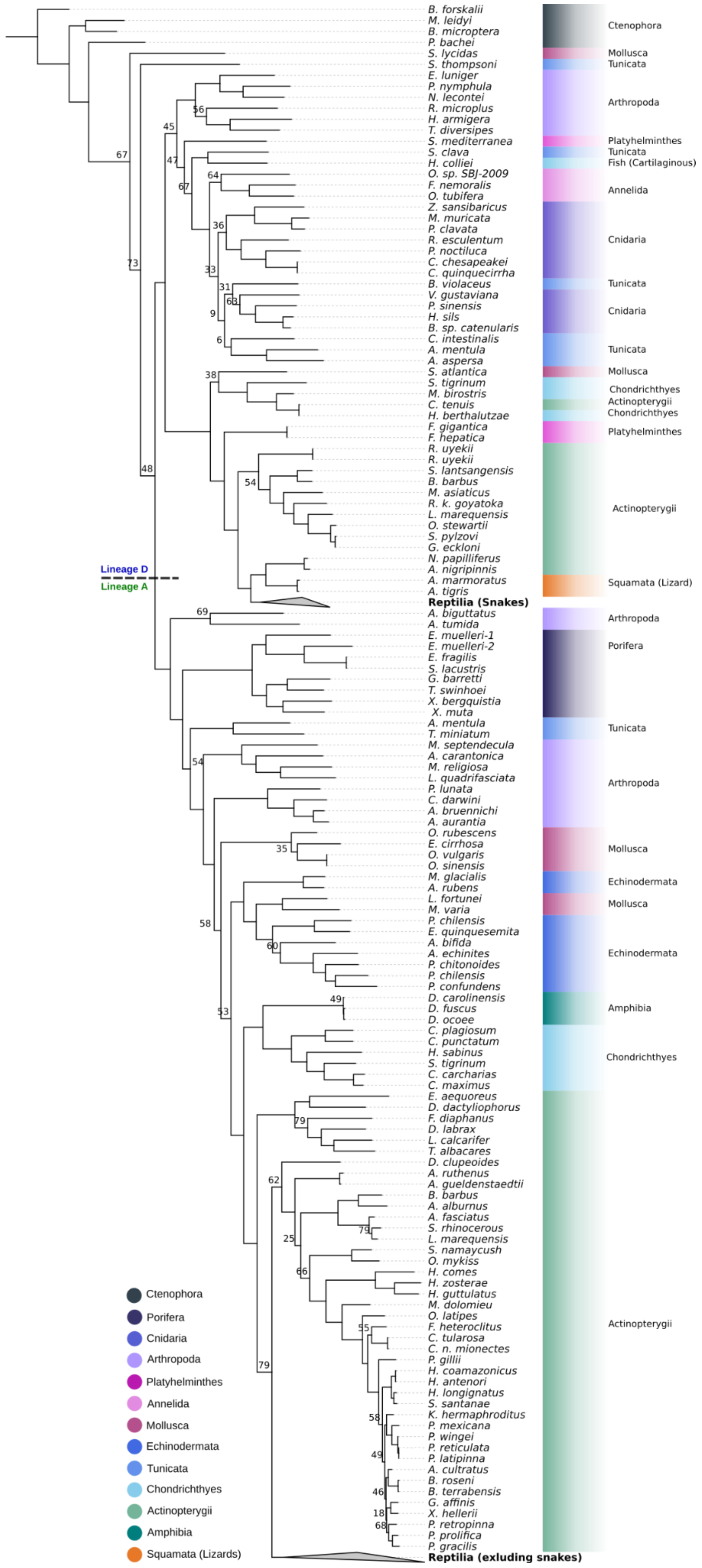


**Figure S4:** Phylogenetic tree of R2s constructed with RaxML (R2 trimmed amino acid sequence, JTT, 1000 replicates). Bootstrap values under 80 are shown at nodes.

**Supplementary Figure 5**

```
R2-1_BMi    VDRVTIDAESGLPNHAGANLLQCEWCDRLCKNKAGLTLHKRACKNNPAVGSSAGNTNDNR
R2-1_MLe    ITRVRTSSNRG-EHSNGVTYPRCE------QGVAPLDTHGGICDAPPQVTVPATETDKQ-
R2-1_PBa    --------------------HRCPNCRKLCRSGNGLALHMKHC-------AKCYQNGDNR
R2-1_STig   MPEATASAH---SSQRGTAMFVCEHCGKEYRSRSGLSGHRRMH-------FAEGETRAV-
R2-1_MBi    ------------ESQIG---YVCPDCGRAFRSRSGMSNHRRTH-------AAAGAVGGR-
R2-1_HBer   ------------KSQTG---FPCPDCGRVYRSRSGMSNHRRTH-------LAQEGGDDG-
R2-1_CTe    ------------KSQTG---FPCPDCGRVYRSRSGMSNHRRTH-------LAQEGGDDG-
R2-1_ATu    ------------DNSTLNIDRRCGLCGVTFNTKSKLKSH-----------ILSRSCSDR-
                                               :  *

R2-1_BMi    RINTPP---TMRSLFNCEYCNTGYGTDRGLSAHISKKHIPEWN---------------MI
R2-1_MLe    --------------KKCEYCEFTYLKPRQIGTHMRKRHPQEWN---------------DI
R2-1_PBa    QEPVKP-----R--MECSICGLFFSGQRGVAIHKRKKHPAEWN---------------ET
R2-1_STig   --PI----------WKCDICQAAFGTKIGLSQHQRRQHAEVDN---------------QR
R2-1_MBi    --------------FSCGICSESFLSKAGLAQHTRHRHPVEHN---------------IR
R2-1_HBer   --------------FRCDICSAVFRTKAGLGQHTRRQHPVQHN---------------IR
R2-1_CTe    --------------FRCDICSAVFRTKAGLGQHTRRQHPVQHN---------------IR
R2-1_ATu    --------------SACKFCGRSFNTFAGVRQHERRVHPLEYASDLQSVIGKASESVIME
                            *  *   :     :  *  : *
```

**Figure S5:** MSA of R2 ORFs with two ZnF N-terminal architecture: Ctenophora: *Bolinopsis microptera*, *Pleurobrachia bachei*, *Mnemiopsis leidyi* (re-curated, first found Kojima 2016). Chondrichthyes: *Mobula birostris, Stegostoma tigrinum, Hypanus berthalutzae*. Actinopterygii: *Cynodonichthys tenuis.* R2-1_MLe has two ZnFs, but the first has a small deletion. CXXC residues in the ZnFs are highlighted in green.

#### Supplementary Figure 6.1

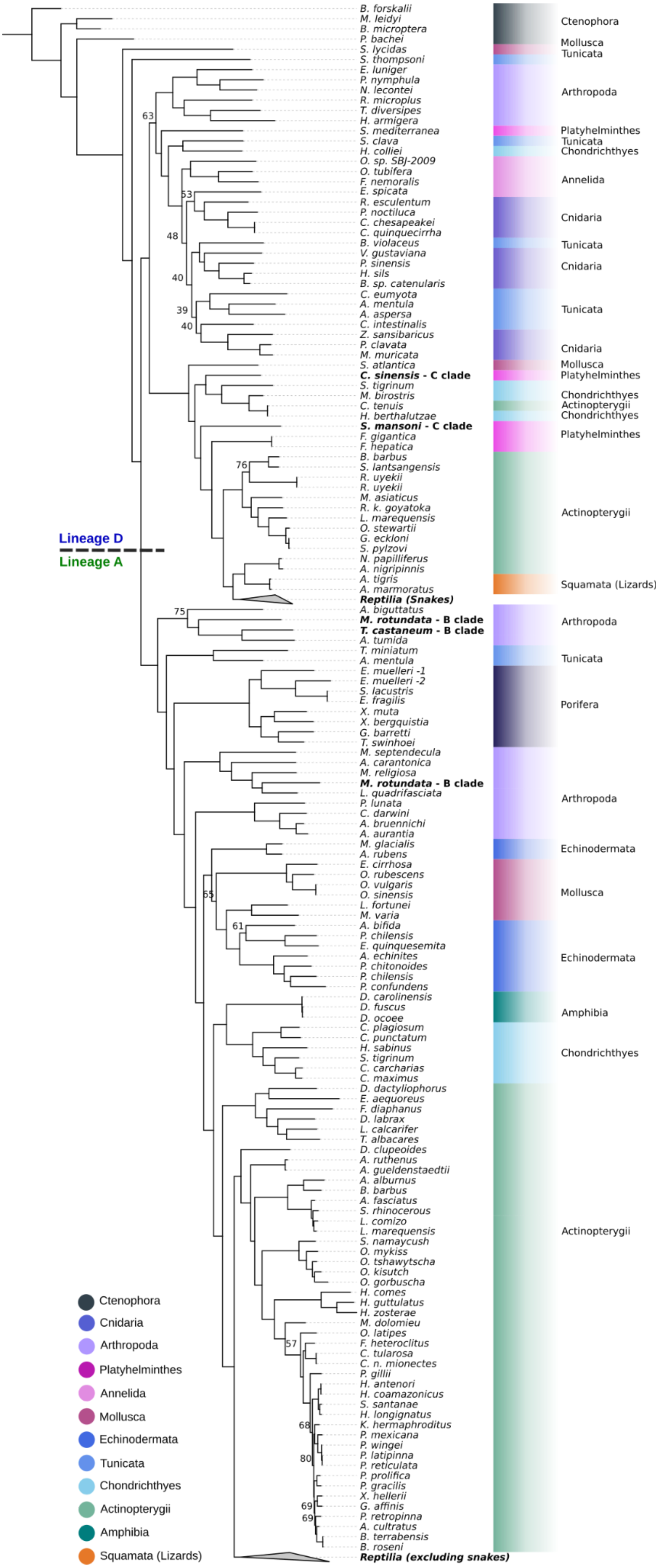

**Supplementary Figure 6.2**

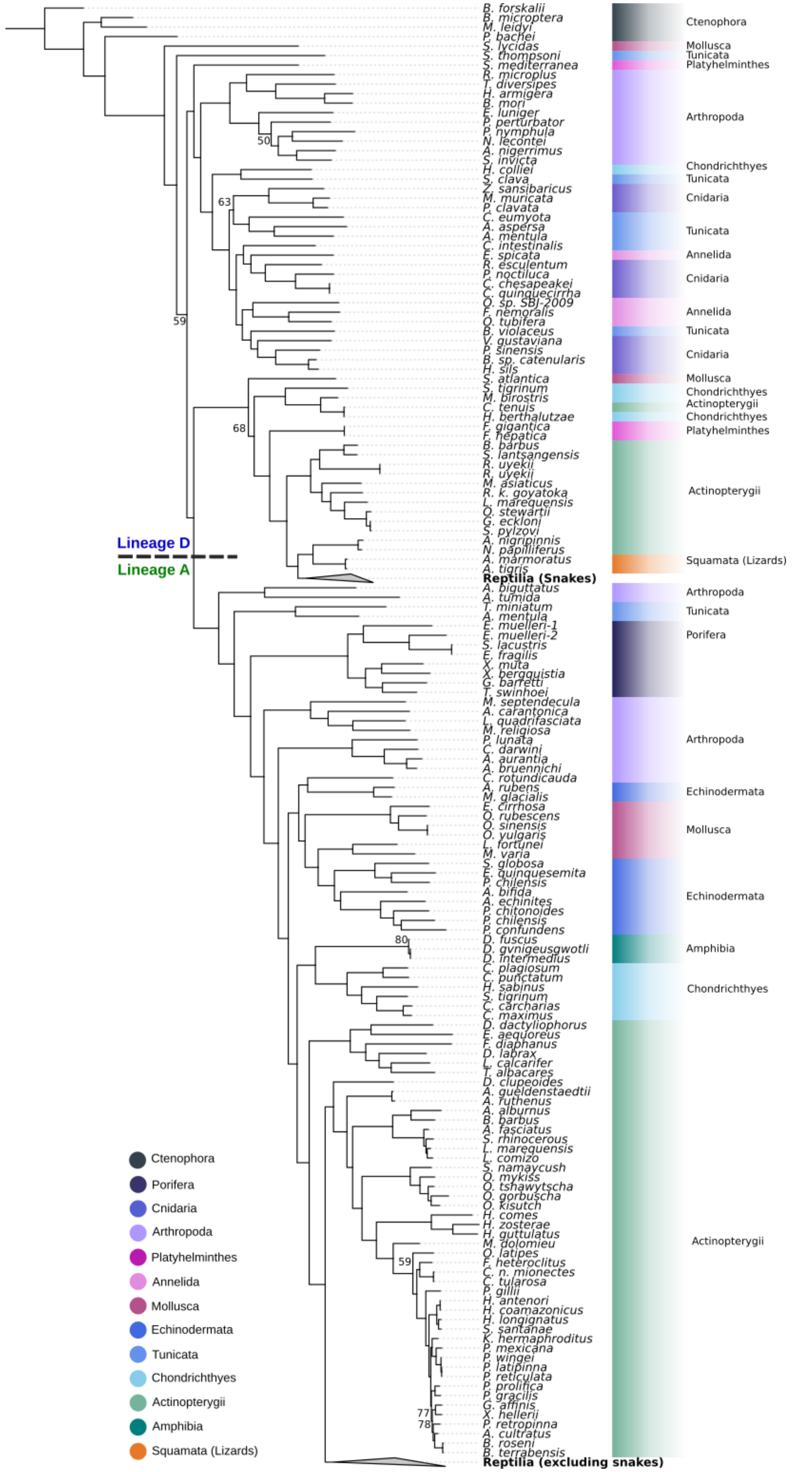

#### Supplementary Figure 6.3

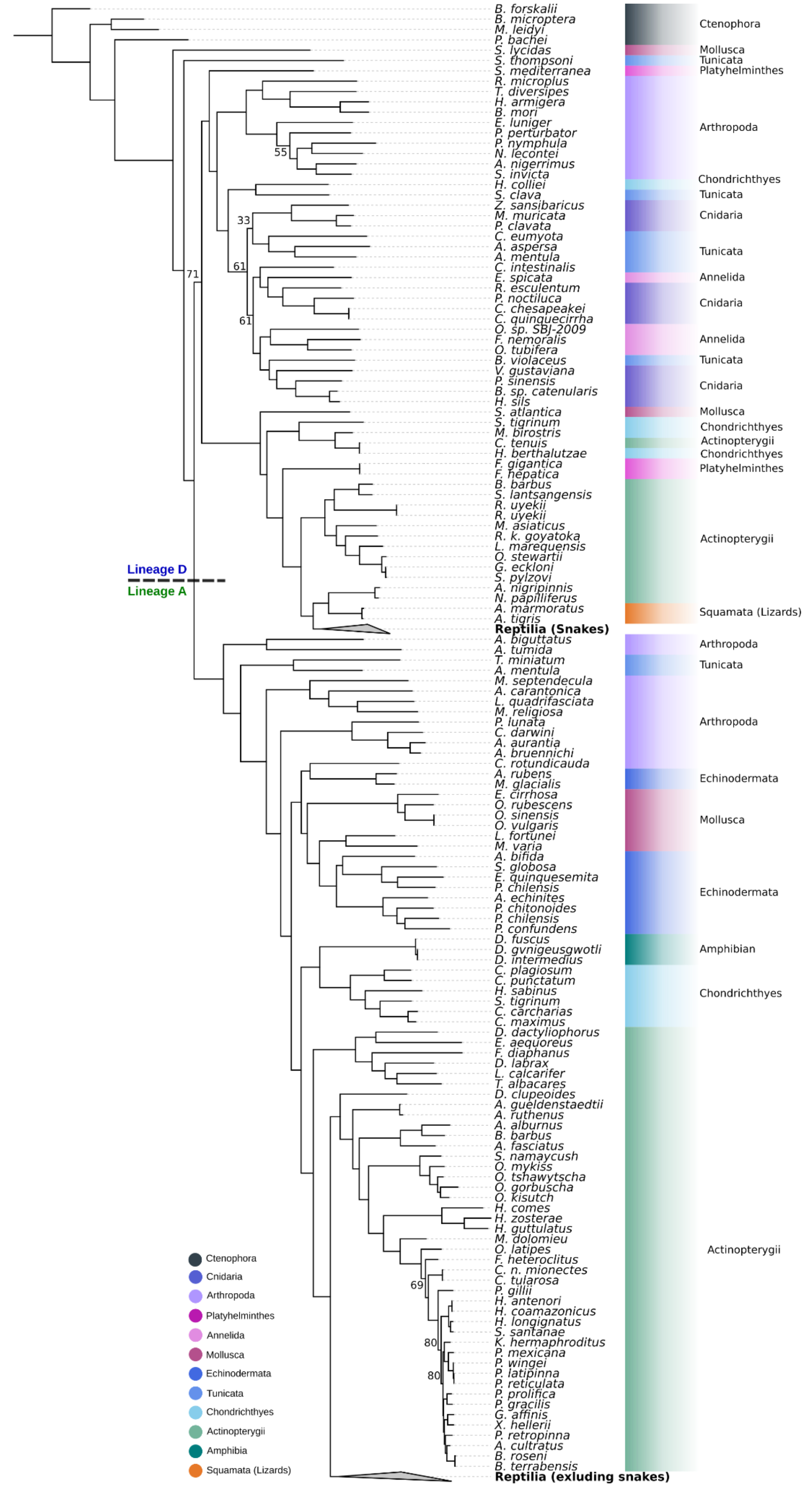

**Figure S6:** Phylogenetic trees of R2s constructed with IQTree (R2 ORF, MFP model finder, 1000 replicates). Phylogenetic trees are rooted with Ctenophore R2 from *B. forskalli*. Bootstrap values under 80 are shown at nodes. (1) Phylogenetic tree includes R2s with two N-terminus ZnF R2s in bold text from [3]. *S. tigrinum*, *M. birostris, H. berthaluszae* Chondrichthyes also have 2 N-terminus ZnFs. (2) Phylogenetic tree of R2s with R2s from reptiles in lineage A and snakes in lineage D collapsed. (3) Phylogenetic tree of R2s with Porifera excluded from tree construction. R2s from reptiles in lineage A and snakes in lineage D are collapsed

#### Supplementary Figure 7

```
R2-1_BMi    ----------------------------------------KNTMFNITPRPEDSQVDRV
R2-1_MLe    KLHRCCPLFQEKDSGGLNRSASGVMSNTSHSKLNLKMDNKLKTSLETPSGVRADSIITRV
R2-1_PBa    TL----PFLKLNMNRLNNEKNSGAVRSTEFS----VADNRPSQSLRTTESH---------
                                                        . :: .
R2-1_BMi    TIDAESGLPNHAGANLLQCEWCDRLCKNKAGLTLHKRACKNNPAVGSSAGNTNDNRRINT
R2-1_MLe    RTSSNRG-EHSNGVTYPRCE------QGVAPLDTHGGICDAPPQVTVPATETDKQKK---
R2-1_PBa    -----------------RCPNCRKLCRSGNGLALHMKHC-------AKCYQNGDNRQEPV
                             :*        :.   *  *   *          . :...:::
R2-1_BMi    PPTMRSLFNCEYCNTGYGTDRGLSAHISKKHIPEWNMIKMERFKADGPRQHVWREGDWEI
R2-1_MLe    ---------CEYCEFTYLKPRQIGTHMRKRHPQEWNDIKRTKFLSE-KRQKRWLDEDFEL
R2-1_PBa    KPRME----CSICGLFFSGQRGVAIHKRKKHPAEWNETKRVEDTLS-RKKIRWTSGDREL
                     *. *    :    * :. *  *:*  ***  *   .    .  ::  * . * *:
R2-1_BMi    LCQGEEEHDRLPSTAKNKLGVNMFIRASYFPQLTIQAISCQRKSPNFHRY--KNDRNLQD
R2-1_MLe    LCIGQEEYLVLSSIGKQGKGINQYIQTKYFPTLSTDAIKSQRKSRRFSEYSEKRSRELQP
R2-1_PBa    LHLGMIEWEASAKTHSQ----EDWVRDHKFPERTINSIKCQIRSKLFRRYVAARPRDEAP
            *  *  *     ..  .:     : :::    **  : ::*..* :*  * .*    . *:
R2-1_BMi    PVTEPQPELEPIPEIEIPAQASNPTNNPLEFTPLAEDIADEIRGKPLGPEILNVYRLLAQ
R2-1_MLe    CNTSSDPE-------ELPNEAVT-ENSPLSFDPLDRDVVKKISSKDHGDQILLVQEHLIN
R2-1_PBa    ---APTPE-AAIQEDEC-------AGSILC---LKDAAIRWLEKEPDGEEYLAIYERLQD
                . **        *           .. *     *         :  :  * : * : . * :
R2-1_BMi    AQFNEANDKSRFIFDELAKDFTRLSTVPKQPTTSKNKNKKKVGKKRISQRPEKKRLSPSK
R2-1_MLe    GRYQEANTLAKAIFEKLSGKFPNLKTGDHRP--GKQQTARKVGKKRV--RGSGKKLSPSK
R2-1_PBa    GRLGEAGERSRRKFDELASKYT--------PLAKSQKPAKGRSKPVDLKSKTNGKLSPAK
            .:  **.  ::  *::*: .:.          *    .::  :  .*            :***:*
R2-1_BMi    AKRKELYAIVQKKWH-TKKRSSVIDTILRDSLGKV---REPAHTPEQLAEFWKGLFSRES
R2-1_MLe    QNRRELYAIVQKQWR-TKKRSKVINQILTGNLNKE---QSYTHTPDQLAQFWSTLFGRVS
R2-1_PBa    RRRRAKYGQCQKEWHNNNQRSGLIRSILQGKFGSKEVLKDPRETEETI-DFWSGIFNRDS
              .*:  *.   **:*: .::** :*  **  ..:..     :.   .* : : :**. :*.* *
R2-1_BMi    PPDNRPIPNPRTEIPQLDNPILVSEVTDSLKRATEKATGLDGVPLKHLREIGATALTILY
R2-1_MLe    PRDDRPINHRRSVIPELDKPLSVEEVEAALKGAKDAATGIDGVPISHLKHLGSAALTILY
R2-1_PBa    TPDSREIENARETIGALDNMITVEEVAKALKGKSERARGPDGVPHKCLKELGATKLAILY
            . *.* * : *   *   **: : *.**   :**   .: * * ****  . *:.:*:: *:***
R2-1_BMi    NGLYCKQLIPTSWKEARTVLIPKCDVPSSPGEYRPITISSYYYRIYTSTIGRRLSDSVGL
R2-1_MLe    NGLYVTGSIPDPWKRARTILIPKSNPPASPGDYRPISISSYFYRIYTSAISKRLASAVSL
R2-1_PBa    NGVFCTSEVPDSWREARTVLIPKKEEPKGPADYRPITIGSYFYRAYTSVLGGRISDVVRP
            **:: .   :* .*:.***:**** : * .*.:****:*.**:** ***.:. *::. *
R2-1_BMi    SNRQKGFIKADGIRDNLILLETIIEDSKKTSSPLHMTFMDVKKAFDSVSHHSIRRALEWA
R2-1_MLe    DDRQKGFIKEDGIRDNLSLIDTLINETKAGSKSLFMTFMDVKKAFDSVSHYAIARSLEWA
R2-1_PBa    SQRQKGFVRSDGIRDNLCLLDGLIYNSKNRVQPLHMSFMDVRKAFDSVSHFSIQRMLRWS
            .:*****:: ******* *:: :* ::*    ..*.*:****:********.:* * *.*:

R2-1_BMi    GVPLGLRNVIADLYKDCSTRIGLSDIKVTRGVKQGDPLSSILFNLVVEMALSKIPDRLGI
R2-1_MLe    GVPDGMRSVIADLYQDCTTDICGRSVKVTRGVKQGDPLSSTLFNLVIEMVMSNVPERLGI
R2-1_PBa    GIPPLLRSVINDLYIGATTSVLGKTVPVKVGVKQGDPLSSILFNLVLDMALDGLTDELGV
            *:*   :*.** *** ..:* :      : *. ********** *****::*.:. :.:.**:

R2-1_BMi    GYLGHRLFYMAFADDLVILSRSQLTNQTLVDRVTEQLGLVGLELHPNKCKSVAVRADAKR
R2-1_MLe    QFQGHRLFYLAFADDLVLLTRGPTANQKLVSLVHEQLARVGLELHPGKCKSIAIMADPKR
R2-1_PBa    SYLGERLCWMAFADDLVILAPSRATLQELISNITDRLKRVGLVMNGDKCKSLSICADGKR
              : *.** ::*******:*:  .   : * *:.  : ::*   *** :: .****::: ** **
R2-1_BMi    KTTFVDSTQTIRVNGSEIPALNSEGWYKYLGIKVSSSGMPQGGYTDEIKLLLERVTRAPL
R2-1_MLe    KTTFVDQGSSVLIGGEPVSSLGPQEWYKYLGIKLGSGGMPQGIYRDQLADLLAKTDSAPL
R2-1_PBa    KRTYVDTSQKLYIEGNMLDSMSITDYYTYLGIGNGARGVKKENLHSEWKTLLERTDKAPL
            * *:**    ..: : *. : ::.    :*.****  .: *: :     .:     ** :.  ***
R2-1_BMi    KPHQRMFILRTHILPRFNHRMMFEKVACKTLTEIDILVREYVRRWLKLPKDVPVAAFYTD
R2-1_MLe    KPQQRLYILRSHILPKFNHRLMFERVTCQTLEGLDKLIRTHVRKWLKLPKDTPGPAFYAD
R2-1_PBa    KPHQKLYVVKKHAYPTLQHKCSFQYASKKCLTELDKLTRRYVRKWMWLPQDTTIEAFYAS
            **:*::::::.*  * ::*:   *:  .: : *   :* * * :**:*: **:*..  ***:.
R2-1_BMi    VPSGGLGLLSLRTRIPLLKRQRTERMAESSDPIIRLLVHQEPSKTRLTIGKKRCRIFGKN
R2-1_MLe    KGSGGLGLITLRYRVPLLKLRRHKKMADSPDPVIRLIPNAEPTISLLARWTKMCSLYGKQ
R2-1_PBa    VEDGGLQLPSFRYEVPLNKFRRLCKMRTSEDPLVRKLANAEPAKTKIDNAKKLCVVDGNS
              .*** * ::* .:** * :*   :*   * **::* : : **: : :    .* * : *:.
R2-1_BMi    YHSKSQLASITREKFWSTCDGKGLRTVVPINTSKSSFKLLSDDRTSLKAAQYLGAISVRL
R2-1_MLe    YQHKSELSKIIRDKYWTMCDGKGLRTEVPPDTAKKTLSLLFEDRTPLKPGQLIGAIGVRL
R2-1_PBa    IENKSQLKSRVRDAYWKSYDGRGLRTE-PQQVKRGNFGQLQGAVTTLSTRNLVGAWNIRL
              . **:* .   *: :*.   **:****  * :. : .:  *     *.*.. : :** .:**
R2-1_BMi    NCLGTPLRNNRGGMKPAI---HNLCDKCPGQKFASLGHISQTCPATHGLRVKRHDKVVTR
R2-1_MLe    NTLGTPARNNRAKGYSPE---ANICDKCPGNRQATLGHISQTCPATHGRRVKRHDKIVNR
R2-1_PBa    NTTQTPARKNRAGGAGGAGDSASTCDKCPNGRLATLGHISQSCPETHGSRTKRHDRVVDH
            *    ** *:**.             . *****. : *:******:** *** *.****:* :
R2-1_BMi    LAKHFKGKENTLTVLVEPQLKYGNLPMQKPDLVINTGSTVEIIDIQIKADQGIPRDEDID
R2-1_MLe    IAKALKERGSVKNILTEPHLRHDKLPLRKPDLIVHTEKSVEIIDVQVVADQGISRHEDED
R2-1_PBa    LQSALLNAQGVTSVLKEPEIRPKGKSYCKPDLVVAMAERVVVVDVQITSDGGIEDLEGV-
            : . :     .. .:* **.::      .  ****::    . * ::*:*: :* **    *.
R2-1_BMi    ETVKKDKYDTQECRRAAYLALGVTPGSLPCNVNAFTLTWRGNPAPHSYKLARRLGFTSIM
R2-1_MLe    QQKKIVKYDVDGYKRAAYKMLGIDYGSIPCNVSAFTITWRGNLAPHSLKLASRLQFSPVL
R2-1_PBa    AKRKTDKYGSPEILKATLLHLGL-PSDTPISVHAFSITWRGNTLRHSLETASTLGVTHIL
               *  **.      :*:    **:     .. * .* **::*****    ** : *  * .: ::
R2-1_BMi    KYLIADALVDTWGMFVVWNCTS------------
R2-1_MLe    KYIVADSLVDTWGAFLIWGKTS------------
R2-1_PBa    PRVTTDVLVDTYRMFLGWRASGRRCVPNKKSTLK
              : :* ****:  *: *  :.
```

**Figure S7:** MSA of Ctenophora B N-terminal architecture sequences: *Bolinopsis microptera*, *Pleurobrachia bachei*, *Mnemiopsis leidyi* (re-curated, first found Kojima 2016). CXXC residues in the ZnFs are highlighted in green.

**Supplementary Figure 8**

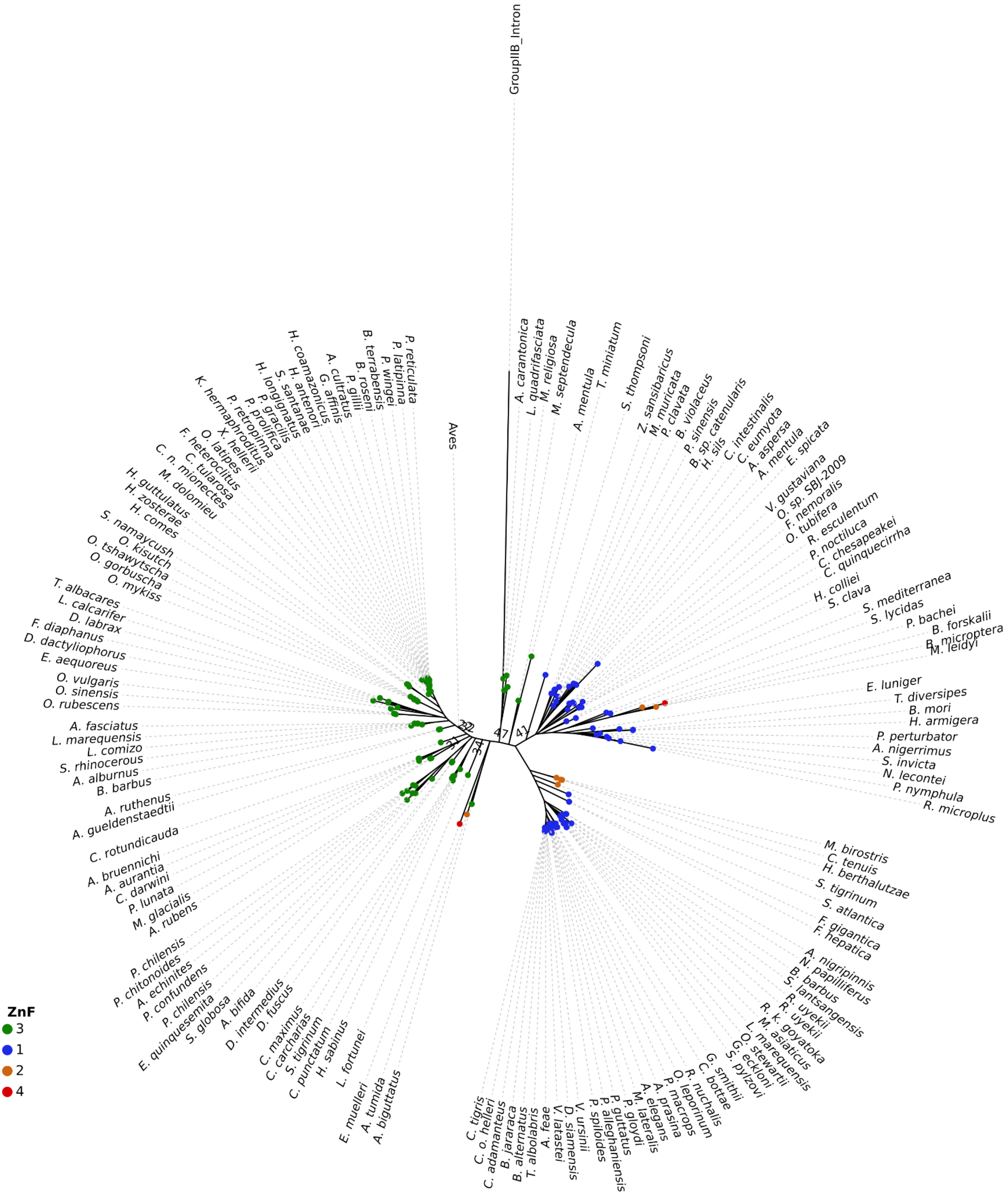


**Figure S8.1:** Phylogenetic tree of only R2 RLE domains constructed with IQTree (MFP model finder, 1000 replicates). The number of N-terminal ZnFs is annotated at each node. Bootstraps under 50 are shown. Aves branch is collapsed.

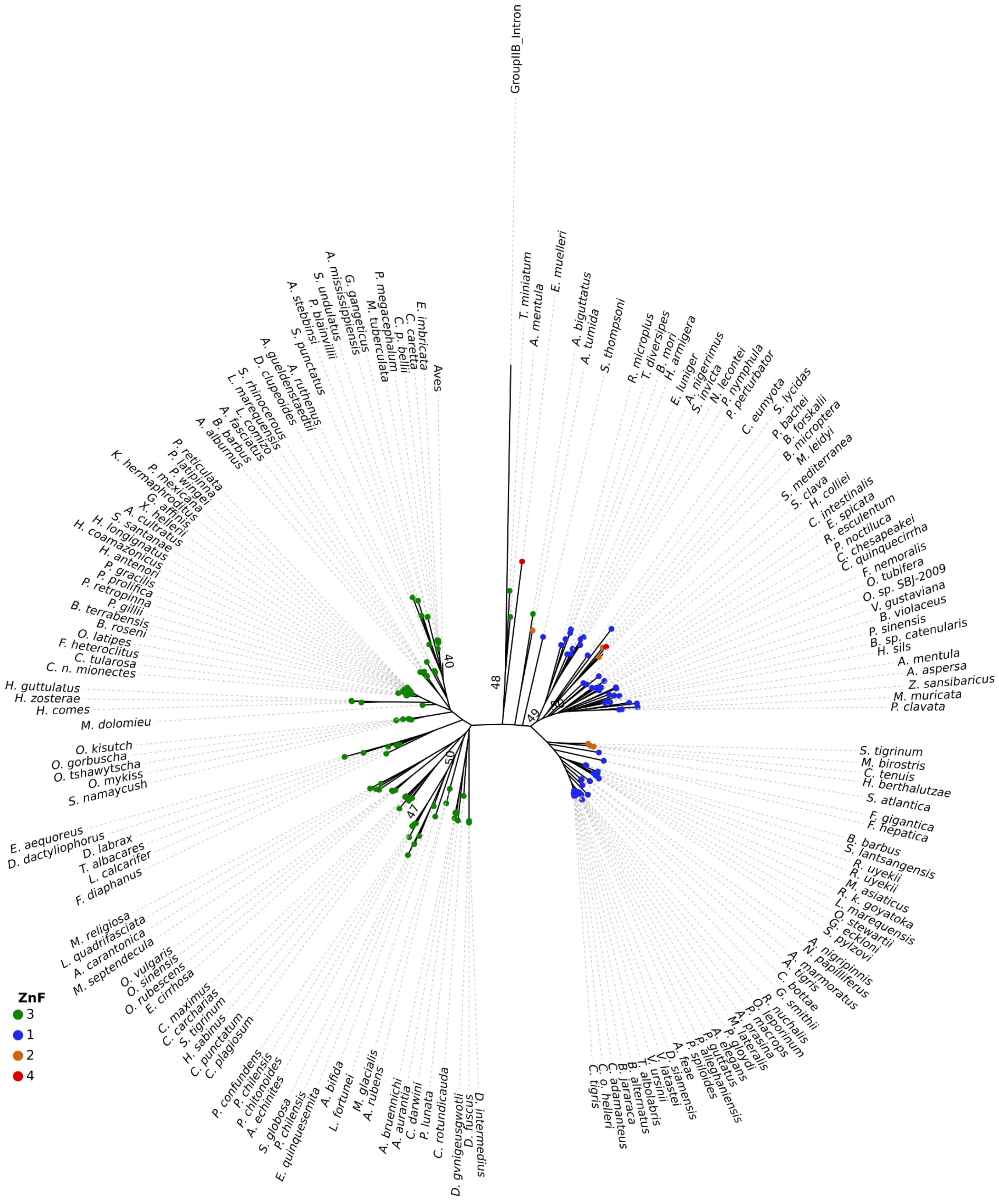


**Figure S8.2:** Phylogenetic tree of only R2 Thumb and ZnK domains constructed with IQTree (MFP model finder, 1000 replicates). The number of N-terminal ZnFs is annotated at each node. Bootstraps under 50 are shown. Aves branch is collapsed.

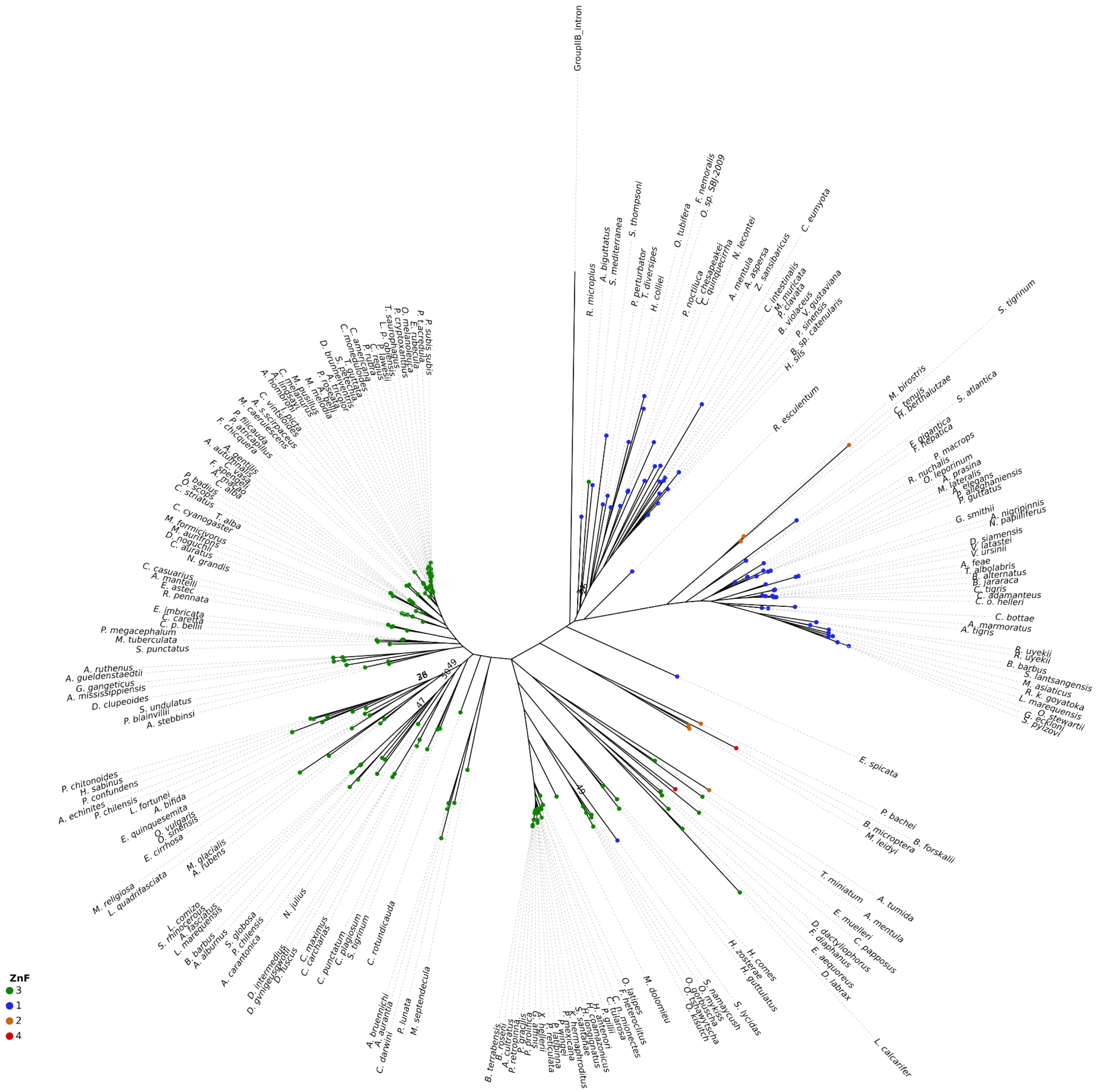


**Figure S8.3:** Phylogenetic tree of only R2 ZnF-Myb domains constructed with IQTree (MFP model finder, 1000 replicates). The number of N-terminal ZnFs is annotated at each node. Bootstraps under 50 are shown. Aves branch is collapsed.

**Supplementary Figure 9**

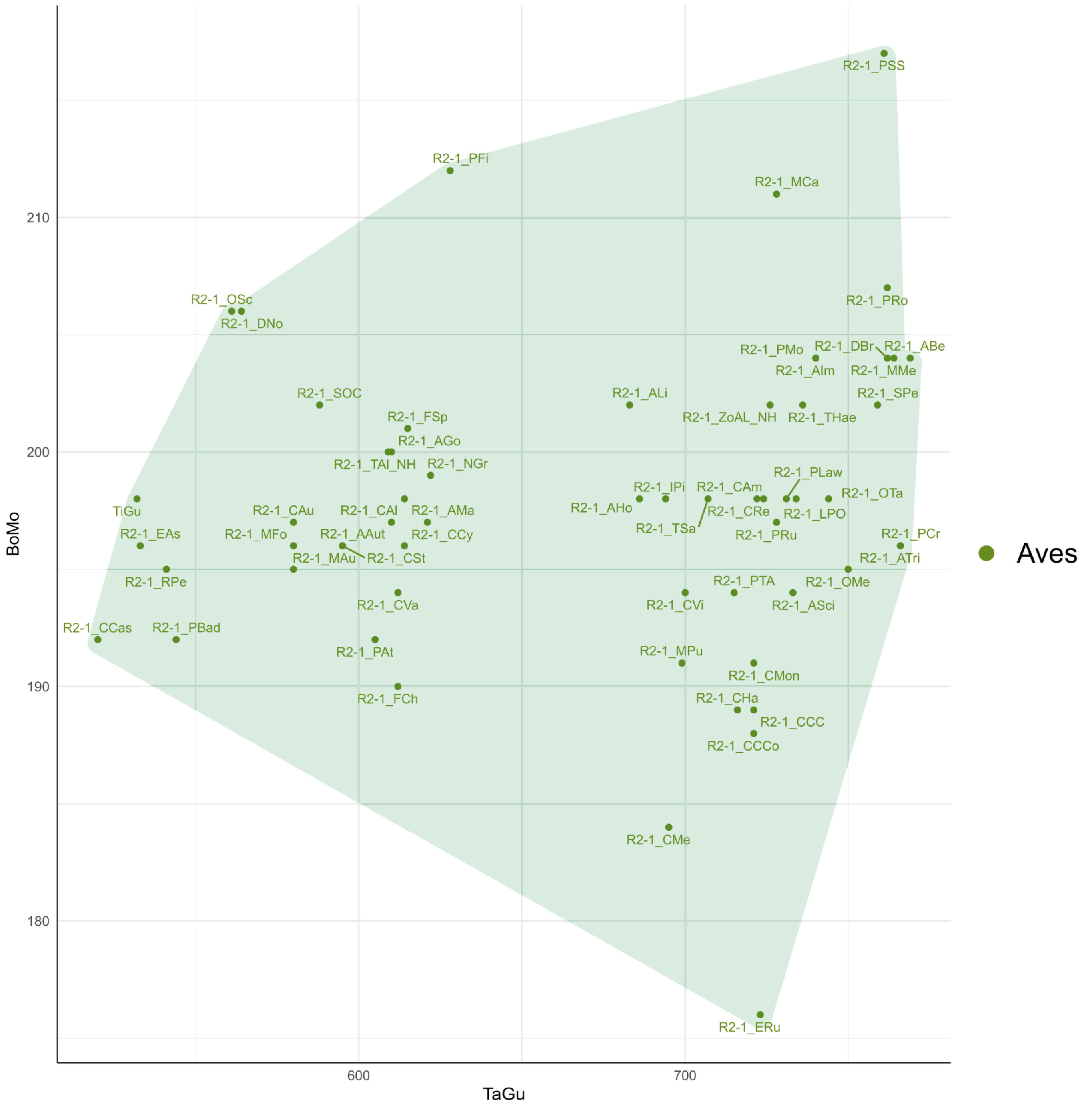


**Figure S9.1:** Position-specific scoring matrix scores of Aves R2 RTs compared to BoMo (*B. mori*) (lineage D R2) and TaGu (*T. gutatta*) (lineage A R2) RTs. Green shading indicates the R2 has 3 N-terminal ZnFs.

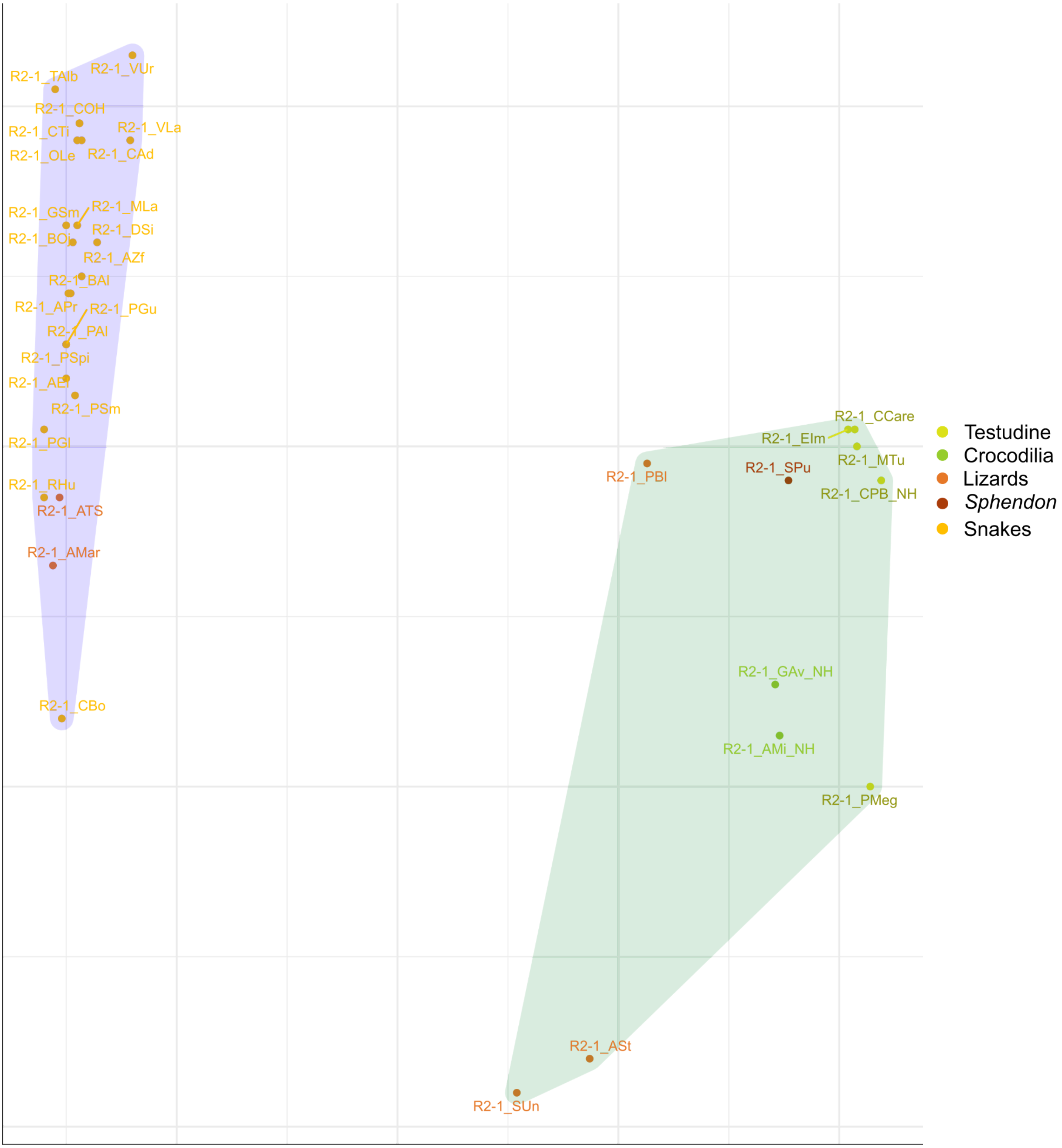


**Figure S9.2:** Position-specific scoring matrix scores of reptilian (excluding aves) R2 RTs compared to BoMo (*B. mori*) (lineage D R2) and TaGu (*T. gutatta*) (lineage A R2) RTs. Green shading indicates the R2 has 3 N-terminal ZnFs, blue has 1 N-terminal ZnF.

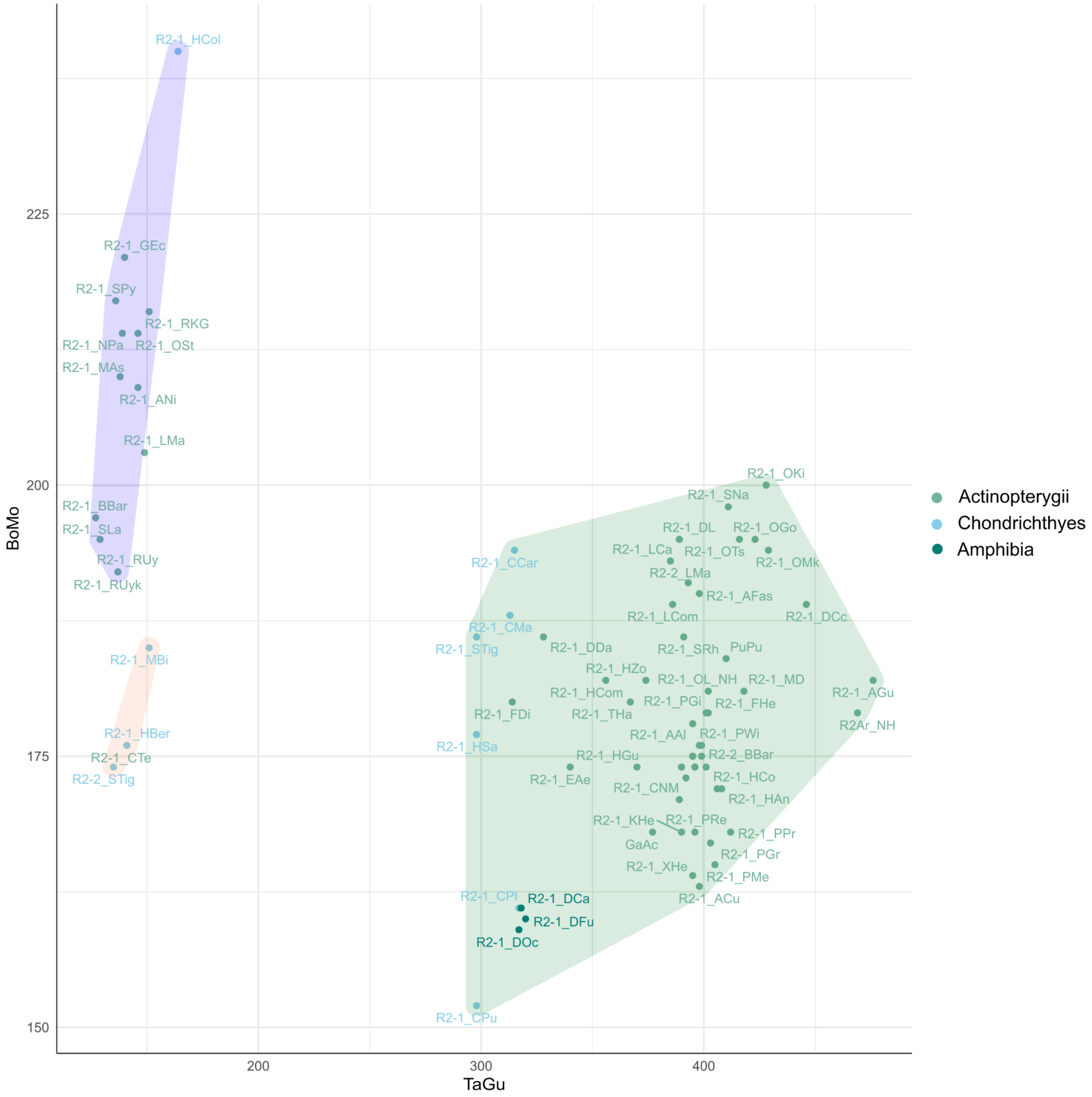


**Figure S9.3:** Position-specific scoring matrix scores of Actinopterygii, Chondrichthyes, and Amphibia R2 RTs compared to BoMo (*B. mori*) (lineage D R2) and TaGu (*T. gutatta*) (lineage A R2) RTs. Green shading indicates the R2 has 3 N-terminal ZnFs, blue has 1 N-terminal ZnF, and orange has 2 N-terminal ZnFs.

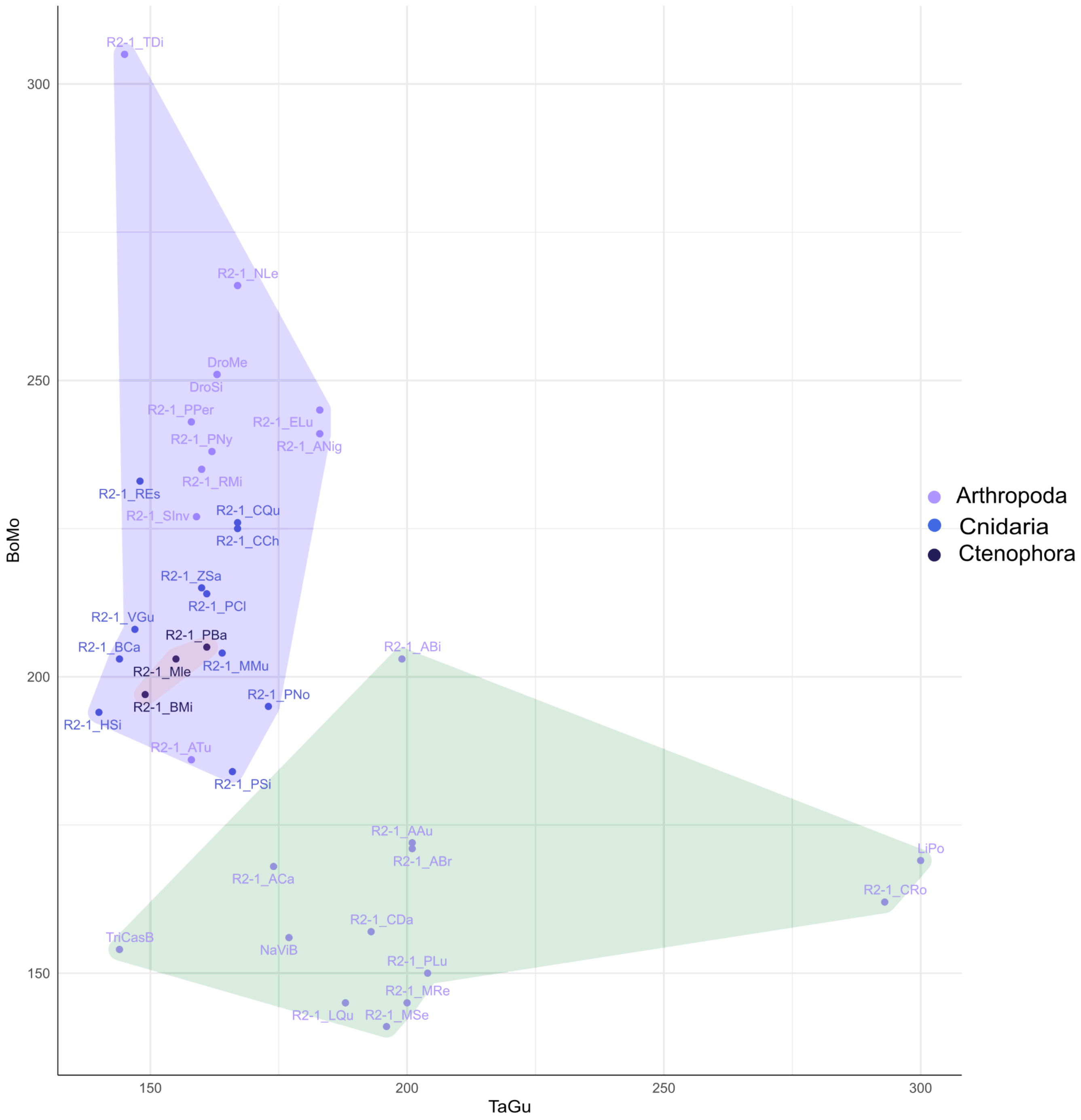


**Figure S9.4:** Position-specific scoring matrix scores of Ctenophora, Cnidaria, and Arthropoda R2 RTs compared to BoMo (*B. mori*) (lineage D R2) and TaGu (*T. gutatta*) (lineage A R2) RTs. Green shading indicates the R2 has 3 N-terminal ZnFs, blue has 1 N-terminal ZnF, and orange has 2 N-terminal ZnFs.

**Supplementary Figure 10**

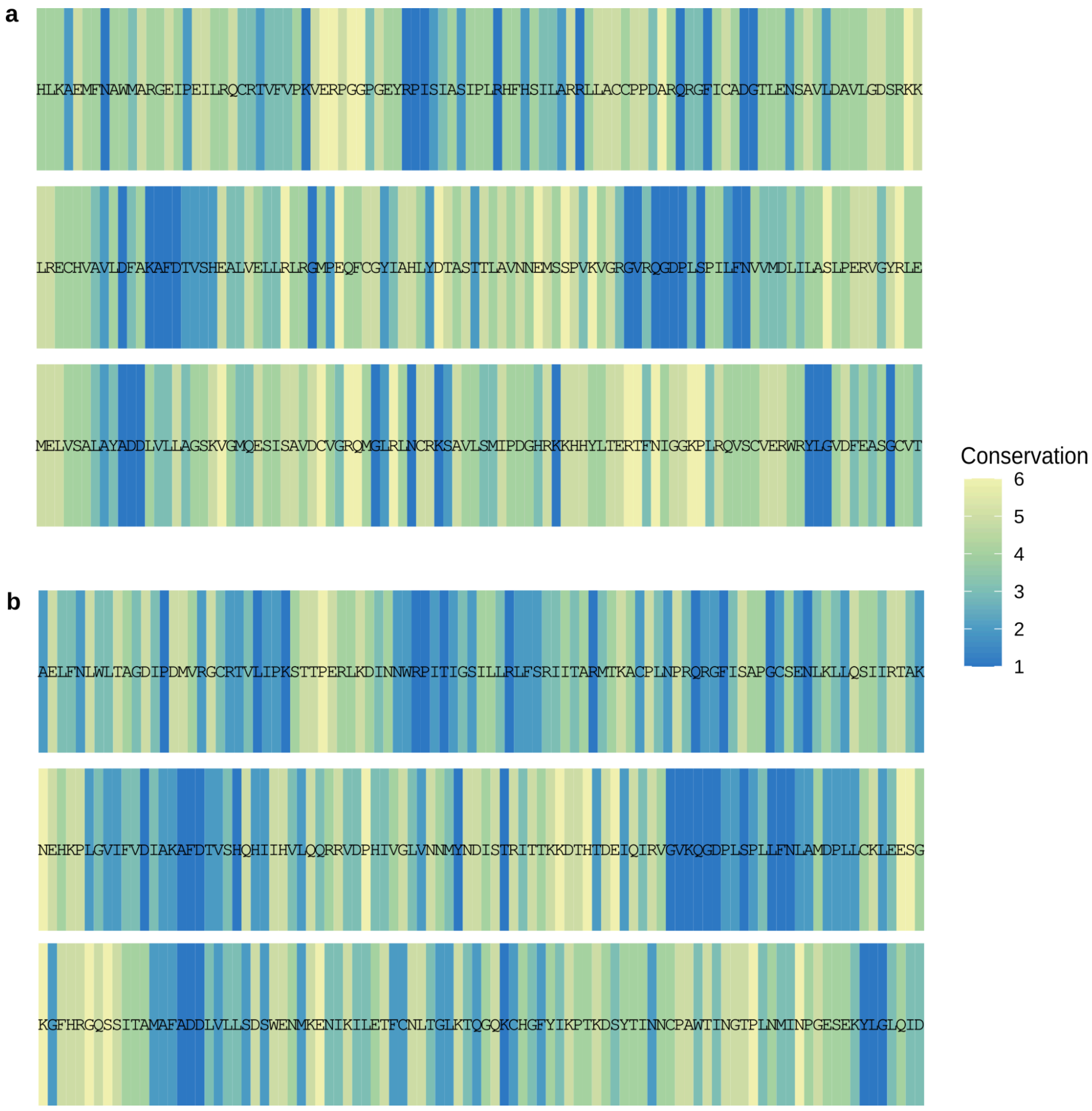


**Figure S10:** Site-specific conservation of individual residues in lineage D and A R2s superimposed with RT residues from (a) *B. mori (*lineage D) and (b) *Z. albicollis* (lineage A, reconstructed from short-reads). Conservation is shown as high (1-3) to low (4-6).

### Supplementary Figure 11

```
LaMa      ------------------------------------------------------------
NePa      ------------------------------------------------------------
CroHe     ------------------------------------------------------------
DeFu      EPSSSKKHPVH-TNATMALPD--AYEVPREQKQLKADIKKKIIYWSYPSPFRCNLCGDIM
AgTri     --------------AGEGTPEALPPVPGENGGQSPDNIVKVTVPDKNPP---CPCCGTRV
GaGa      ----PESAPVQRTEADTTLPS--ASAPAGEGEPPSTRVFLVRLPDSNPP---CPICGDHV
SPun      RPIHPEEESAICVTANESTPQIEYNIIERETDVIPSSSAIVQLPQTNPA---CPFCADRI
PlaMe     EPI--------------------INGTQQK--------TIIQLPNDNPA---CPFCGDHV
PBla      ----------------------------KRERKLDKDEPTIWLPDNKPE---CPYCKEVV
                                                            ****

LaMa      -------------------------------------------------------DNITC
NePa      -------------------------------------------------------ERVPC
CroHe     -------------------------------------------------------PKHSC
DeFu      STIKELKRHISCKHKTFNLVVFCSKCDKEDA-FHNIACHFAKCNKKIIED-----ATFMC
AgTri     NSVVSLLEHLKGSHGKKRVRFRCAKCGKENVNYHSVICHFPKCKGAEIEK--VPAGEWIC
GaGa      GKPSALAVHLVESHAWADVQYQCTHCEKVSSNKHSILCHIPRCQGRVTDR---SDRNWAC
SPun      TTPALLKKHVKGYHGGKLLQFQCSKCGRLNENQHSIITHLPKCKGNSIDP---ALGEFHC
PlaMe     GKPSALNVHLKRNHGGREVEFQCSMCNKADPKAHSILCHIPKCKGKVTEE---PTGDWAC
PBla      GKPTALQKHLTGVHGNQAVTFGCMKCQRKDPRVHAILVHIPKCKGPNIPDARQAQNKHRC
                              ****                                 *

LaMa      RDCGRAFPSLRSLGMHRHHQHPEEVNEERLELIKKK--------RKQWSKEEDEQLMSTA
NePa      -FCGLTFATKRGLSMHQHHRHPTELNAARLTALPSR--------RGVWTSGEDEALIRLA
CroHe     -PCGSEFGTKRGLAMHRFHCHPAEVNAERLTTLPTK--------KLVWTQEEMDGLLRHA
DeFu      RHCDERFTTKSGVSQHARHRHPKAVNDIRLKIAMPA------KRSKLWTLEETLLLMDLE
AgTri     EVCGRDFTTKIGLGQHKRLAHPLIRNQERIVASQPKETSNRGAHKRCWTKEEEELLIKLE
GaGa      VQCPASFNKKVGLSQHKRHVHPVTRNAERVAGSLSRAGLRPRTRRGCWSVEEETLTRLD
SPun      EICNNNFLTKSGLSQHKRIRHPLVRNEERIMASQPKENSLRGKHKSCWSNDEVEQLKTLF
PlaMe     ETCNKQFNTKSGLSQHKRIAHPAIRNQERIAASQPKPNSQRGKHNSCWTVEEEQLLAAFN
PBla      EDCGNQYDTQSGLSQHTRHKHPETRNKQRIEEKEKEKGGKRGTHKSCWTQEETEQLALLW
          ***    : .   .:. *     **   *  *:                 *:  *   *

LaMa      DMEWKVGMMKKDHLAAIRIKFSHRTLDSIKKRLQHLGWAPPPAGQQPQNTPLPARSRPPR
NePa      NDIWPSVTLKKHLYNALMPHFPGRSAEALKKRLQLLQWLP--------------------
CroHe     NGAWKEGMLKTQIFTSLMPHFPGRSAEAIKKRLQSLKWVP-------------ARDNPER
DeFu      ATYKN----SKKINMDIQKHFENKTSIQIGEKRRELREKT-------------AKMEKEA
AgTri     AQFEG----NKNINKLIAEHITTKTAKQISDKRRLLPKKL-------------PEVTDKE
GaGa      ALFRG----ARNINQLIAAEMGTKLPKQISDKRRQLGLCP-------------EQATSGG
SPun      DEFGK----YKDVNKRIANLLPSKTAKQIADKRRLLNLYI-------------KSTKTPN
PlaMe     NMFWG----KKNINILISDHIHMKTAKQISEKRRLLGLNK-------------NATVTTT
PBla      EKYEG----HANINKLIQQELHTKTAKQIGEKRRNILEKR-------------NNKPEIG
                    .    :   :  :    : .: : :

LaMa      RIARASSPPLT---SALAGPLPPSPIGSPPPQAKRRGALWTNREDSELEEHARRLWKPNM
NePa      --------PL----TPSHETLNASAL---SP-----------------------------
CroHe     -----NSQPL----HATSGNVAPATLPTPDP-----------------------------
DeFu      -----NL-------HIDSDDEIESINKSEDE-----------------------------
AgTri     -----PGVCLRTRRAAGSVGTEPGTSQQSQA-----------------------------
GaGa      -----DA-------ESTSVVEEESVTPEVET-----------------------------
SPun      -----PI-------EPET--QEVTIPENRES-----------------------------
PlaMe     -----NP-------LPVS----STCHLKIRT-----------------------------
PBla      -----EE-------IPKT----KTQSINRQT-----------------------------

LaMa      LKKSLAEALSCIIKGRSAEAIRKRLQKLGISSATMTSREDIDSEAMRDKDIHQTTPCVQP
NePa      ------------------------------------KALD--------------------
CroHe     ------------------------------------SRVD--------------------
DeFu      ------------------------------------EHTN--------------------
AgTri     ------------------------------------GILD--------------------
GaGa      ------------------------------------QSPL--------------------
SPun      ------------------------------------SAPV--------------------
PlaMe     ------------------------------------DSPN--------------------
PBla      ------------------------------------------------------------

LaMa      EEDGGYETPTSKKEEWWRTLLTKALETL-NEARIESDKL------------------KL
NePa      ED-------LI---RWKGRMLDAISPNL-SEPRLGTEEL------------------LS
CroHe     DQGANIEDPAA---SWRRALLAAAADHF-KEPKLGAGTL------------------KQ
DeFu      VPQSDVQI------RNTRISTHKKSPQI-KDHKDLVERITEIKDCMKNPEYYDCGIGLME
AgTri     NGPGKCHL------PGRPAAGEKTMEKL-RRHPDKDNGR------------------QK
GaGa      KPPGKI----------RKVLAQRARWL-KKGQDLSDKV-------------------RE
SPun      --KGLC-----------EALQSKARKNLEKESAMKFE---------------------KD
PlaMe     TTTGLK-----------DTYMCKINENIVNQGQIKFD---------------------SE
PBla      EKEGLA-----------NKFKQTAITLI-QEHRIKNESI------------------AR
                                       :

LaMa      MAKGMLEG---TMTREEAAEQLECIT----------------------------------
NePa      LVCNLRDG---SLSSEELNARLEAHA----------------------------------
CroHe     ITQDLLNG---KISSGECRVLMGEHA----------------------------------
DeFu      VITQYIT-----ENISTIKPSASVRPTTDDPAAPNDATE-IEV-----------------
AgTri     TSGQRREGLLQAHYQKTIKEGLSAGAINNFPGAFKQLMDGREMRTMINQTAQDCFGCLES
GaGa      VLGAWVEG---QPGIRARVESVSLDLVLTSFLGAPSG------------------------
SPun      FLTNWLDN---MQNVRHLLEETTVDALCNFLPEPKN------------------------
PlaMe     VISAWMAG---DSNIRSLVESTSLDILSTFLMETPK------------------------
PBla      ALHTWLQE---NNNTRQTVEKATEEILNNFPRTAG-------------------------
```

```
LaMa       -------QEFAPLKWKPREKRTGHR--KEPRSNKQIRRARYAHIQRLYKLKRKDAAHSIL
NePa       -------ARHFPHLWRPSTKRTQSI--NKP-SARQIRRANYAAIQRLYQTQRKDAASSVL
CroHe      -------QHWFPHTWKPSSARPEPK--ARQWKTREIRRANYAAVQKLYKLRRRDAAQAVF
DeFu       -------PTNVPQNAEPTTASSSHKRNSGHKKRYAKRKGSYKEYQTMFLKDKKKIARIVC
AgTri      ISQIRTAMRGKNSGKKATMEQPARKF-QKWMKDRAIKKGNFLRFQRLFHLDRGKLAKIIL
GaGa       -------PRGAPDKKRP--EKGGST--TSWMSRRAVKRGVFLKYQRLFGTKRKLLADIIL
SPun       -------ITKPKKDRRP-VKDRKQA--KSWMKKRAKKRGLYQHYQSLFLKNKTRLASIIL
PlaMe      -------PRKKGNNKIT-NKKSGKK--KKWMEKRAVKKGFYKRYQHLFETDRCKLASIIL
PBla       -------IHNIPKRKKTKKPRGPYR--GKWRQKRRIQKLTYKQLQELYDKDRAAAAAYIL
                           .              .    ::  :   * ::   :   *  :

LaMa       DGKWREAYREKEREVEGVEEHWAQVFETQAK-DGWPQSM--LPPDLSNIKWGLLEPVTAM
NePa       NGSWRSAYKNRTSITDDLDSYWSNIFLQPSHEDPT------PADPPSTPHWSILDPITAT
CroHe      SGSWRNAYRGRAPQPDKMDQYWAEVFETASATDVT------PPGSPSEIYWSVVGKITPS
DeFu       DGAEDMSCQ---IEATEIFDYYKNILENENTKNPKFELSNIVPSEEDFDHLN--RPITEF
AgTri      DDIECLSCD---ISPSEIYSVFKARWETPGNFAGLGDFK--ITGKADNNAFK--DLITAK
GaGa       DGADRAQCV---LPLEEVLRAYRGKWEVESSFEGLGRFG--VRRDADNFAFK--ALITPE
SPun       DGTEKFECE---IDPKIVYEAYKNKWESATEFKGLSNFH--SFDVTDNSKFY--TRISGK
PlaMe      DGTERLQCQ---IPLTEILETYKSKWETLTPFEGLGQFK--SHAVADNTAFE--ILLSAK
PBla       DGEKKTNCD---LPIQEVYQTYKDKWETKTEFKGLQNFH--PWGGTNNDILS--NLITGE
           ..               :   :                        .         ::

LaMa       EVEVALRSM-NNTSAGMDKLSAQEVLTWDL---PSLAGLLNVILATERLPSTLATARVTL
NePa       EVTSALSSM-KSSATGLDRLSASQLLSWDA---NSIGAYFNILLVSGLVPTHLTMARITY
CroHe      EVANALKGM-RNSAPGLDRITAEELLTWDH---PSLAAYYNLMLAAGGPPEHLACSRVTF
DeFu       EITWHLENSNMNKATGPDNVGLKELFGLHARDHTILTDLFNIWLQTSKVPECIKRNRSIL
AgTri      EIERNVQEMSKGSAPGPDGITLGDIVKMDP-GYSRTAELFNLWLTSGEIPDMVRGCRSVL
GaGa       EVVKHMMAMASKSAPGPDKLTLRDLRRADP-EGDALAELFSLWLITGTVPDGLKECRSVL
SPun       EVQKNIKEMSRKTAPGPDGITVEDLENVDP-DGEILAALFNLWMAVGIIPDTIKECRSLL
PlaMe      EIMKNIKEMNKNSAPGPDKVSLRDLLLADP-ECNALEKLFNTWLITGIIPNSIKECRSLL
PBla       EVKEHLQAMCNKTAAGPDNISVKDLKQVLH-VEQKLAELFNLWLITGQIPNTVKKSRSIL
           *:   :      .:.* * :   ::               .  :     *  :   *

LaMa       IPKVEEPS---GPNDYRPIAISSVIARALHKVLSKRMREQFEFSPLQYAFLQRDGCLEAS
NePa       VPKTDSPS---GPSEYRPISVTSVLLRAMHKILARRMLATLDFSDLQLAFLQRDGTLDAS
CroHe      IPKVENPQ---TPGDYRPISVASTVLRAFHKILAWRLRDNLQLSPFQHGFLQRDGCLEAT
DeFu       IPKNVPLDSLGNIGNWRPITIGTALMKLFTKLLTKRLSTFVSIHERQKGFINARGCLENL
AgTri      IPKSTKPERLKDINNWRPITIGSILLRLFSRIVTARLSKACPLNPRQRGFIRAAGCSENL
GaGa       IPKTVDREKLGQLGNWRPITIGSIVLRLFSRVLTARLAAACPINPRQRGFIAAPGCAENL
SPun       IPKTSDPEKLKDIGNWRPLTIGSIIIRLFSRILTIRLAKACPLNARQRGFIDSPGCSENL
PlaMe      IPKTADPEALKELGNWRPLTIGSIVLRLFSRIITNRLAKACPINARQRGFIATPGCSENL
PBla       IPKTSDETEKKKVGNWRPLTISSVVLRLFSKIMTSRITKACPLNNRQRGFIAASGCSENL
           :**          .::**::: : : : : :::: *:     :   * .*:   *  :

LaMa       ALLHAVLRTAHERTKPLAAAFLDVSKAFDTVSHNAILGAAEKAGTPPPILRYLNQLYKNA
NePa       TILHTILRKVHTELKPLSMMFLDVSKAFDSISHHTLIRVATTSGLPAPLLSYLRHLYQTS
CroHe      ALLHTILRKVHNSRKSCAMLFLDVAKAFDTVSHQTLFRVAVELGLPPPLVNYLKCLYSRS
DeFu       NILKNSIKGARKNKDSLAVIFIDISKAFDSVGHKHILNSLKRLHVPLGYRQLIKDLYTNS
AgTri      KLLQTIIRTAKSEHKPLGVVFVDIAKAFDTVSHQHILHALQQRGVDPHIIGLVNNMYKDI
GaGa       KVLELLLRKRKRDRQPLGVVFVDLARAFDSVSHDHISWVLKAKGVDEHIVNLIEDSYQKV
SPun       KSLQSIIEYSKKEKQQFGVVFIDIAKAFDSVSHDHIIWVLKERKVDEHIKKIIQDSYTKV
PlaMe      KILHTIVKQAKTSKKSLGVVFVDIAKAFDSVSHDHIMWVLQERGLDQHIVNIIEDSYKKI
PBla       WLLHNIIKRAKDKKKELGVVLVDIAKAFDTVSHDHIRWVLRERGMDKHIIQLIMSAYDNA
             *.  :.  :   .  .  ::*:::***::.*. :                :   *

LaMa       KLQL------GSTTTQCSRGVRQGDPISPILFILVMGEVLEEALPD-VGVRW----GERQ
NePa       KIRL------GTHDSSCGRGVRQGDPLSPILFILAIEDILHRVLPE-AGFDL----AATR
CroHe      TVRL------ADKATKCGRGVRQGDPLSPLLFIMVMDDIVRKTLPE-VGFDL----DGQR
DeFu       TTTFKGKNNINTDEIHMKSGVKQGDSLSPLLFNIAMDPLICDLQYKGCGFSFNNIQGTRQ
AgTri      STYVTTKRDTHTDKIQIRVGVKQGDPLSPLLFNLAMDPLLCKLEESGKGFHR----GESS
GaGa       TTRVQVFNG-VTPPISIKTGVKQGDPMSPLLFNIAMDPLIAKLETDGQGVKV----GSAS
SPun       STRLKVSKT-LTDSISLKVGVKQGDPMSPLLFNLAMDPLINALEEQGEGVQV----EDMK
PlaMe      HTRMEVGTE-RTPPIEIKVGVKQGDPMSPLLFNLAIDPLITALEKANTGFSY----GKNK
PBla       TTSIKLKEG-NTPDIHIRSGVKQGDPLSPLLFNLAMDPLITELETEGHGVED----ENWT
              .               **:***.:**:** :.:  ::        *.

LaMa       IDSIAYADDLILLAESPRELQRKLDGVCRGLQKAGMALNNKKSVTMTILKDGRRKTLALA
NePa       IGSIAYADDLILLAERPERLQEKLNILLSAFHDAGLIINSSKSHGLTIAKDGKKKLLVLL
CroHe      VDSLAYADDLVLLAEKSPRLQDKLHLLSEALRKAGMSLNARKSRGLTITKVSKRKQMVIT
DeFu       TTALAFADDIAVLSNSWKGMQANLQIIEKFSKATGLKLNVKKTHGFLISHF-GDKIVVNK
AgTri      ITAMAFADDLVLLSDSWENMQTNIKILETFCDLTGLKTQGEKCHGFYIKPT-KDSYTINN
GaGa       LTTLAFADDLVLLSDSWEGMLKNISILEDFCNLTGLRVQPKKCQGFFLNPT-CDSFTVNN
SPun       LKTMAFADDLVLLSNSFEGMSKNIKILEQFCQTTGLQVQPKKCQGFLITPT-KDSYIINN
PlaMe      ITSLAFADDLVMLSDTWEGMNKNIQILETFCNLSGLKVQAKKCYGFFLSPT-HDSYTINK
PBla       ITTLAFADDIAVMSDSCKEMKQNLQIIEAFCNLTGLKVQTTKSYGFALQPT-KDSYIVNN
             ::*:***: ::::    :  ::  :      :*:  :  *   : :      .

LaMa       PHQYTTDNGQVPCMGLSDSQRYLGIQFT-WKGRVTPK-QTSELGRMLLEITSAPLKPYQR
NePa       PHEYRTGSSTIKPIGPTNVITYLGLRFN-WKGQLLPR-HTATLSTMLAEVSQAPLKPYQR
CroHe      PTTYECEGEPIKPMGTDDSVRYLGLHFN-WKGRIVPK-HTGKLDSLLKELTKAPLKPYQR
DeFu       CKPWLFERSKIEFIHPGESERYLGLNFDPHIGCNTPNALNLKLQSWAKKLDDLPLKPTQK
AgTri      CAAWTINGTPLNMINPGESEKYLGLQFDPWTGLAKTN-LTTKLEFWLERIDQAPLKPLQK
GaGa       CEAWKIAGREITMLGPGESTRYLGLNVGPWVGIDKPD-LGTQLSSWLERIGTAPLKPMQK
SPun       CKKWSIEGTEVNMIQPGQREKYLGAKIDPWTIFAEIN-FEEKIEDWLCKLEVAPLKPSQK
PlaMe      CDAWKIDKDSLNMIQPGESEKYLGLKVDPWIGFSKPV-LAEKLTIWLKRLTEAPLKPSQK
PBla       IKPWTIKDQPIQMVQPAHTTRYLGIQVGPWKGMEKPN-MIADTKKWLQNITKAPLKPTQK
              :      :  :   .   *** ..                     .:   **** *:

LaMa       IELVRDFLVPRLLHELVLGCAHRNTIARMDRMIRRETRTWLRLPKDTSLGFLHSPVKSGG
NePa       LEVLRNYLIPKLTHELVLGRAHRNTLKKIDVLIRAAIRQWLRFPKDTPTAYFHARIQDGG
CroHe      LQLLKFHAVPKFTHELVLGHAHRNTVKKLDCLTRAAVRKWLRLPQDTPLGYLHANVKDGG
DeFu       IKIFCQYIVPKLSYSLEMTGTGANTLVALDQTIKATVKHYLHLPHYINDGLLYSRKKNGG
AgTri      LDILKTYTIPRLTYLADHSEIKAGALEALDQKIRTAVKDWLHLPPCTCDAILYSSTKDGG
GaGa       LSLLVQYAIPRLNYQADYAGIGRVALEALDSMNRRKVKEWFHLPACTSDGLLHSRHRDGG
SPun       LEILNIHTIPRIIYLADHTNCNITKLKNLDNMIRKRLKDWLHLPASTCHGLFYSKNRDGG
PlaMe      LTMLNIYTIPRIIYLADHTDTKKTLLSSLDDNIRTVVKGWLHLPPDTCNGFIYTKTRDGG
PBla       LEILRTYTIPRLIYHNEQTGTGKTKLKELDNLIRAGIKSWLHLAHDTCTNLIYTKTKDGG
           : :.  . :*::              :  :*    :   : ::::.       :::  :.**
```

```
LaMa       LGIPCLGTTVPLLQKRRFEKLLSSDCPIKQTLTELPSFMTTLRRVNL-------------
NePa       LGIPALTTQMPFLQHHRFTKLLSSASPSIQSICQEPAFVQAYARVLT-------------
CroHe      LGIPCFSTSIPLLQKKRFEKIATNPATIFQIMQRQDSFRTQGRRLDK-------------
DeFu       LSIPKLCTSIGINKVNNLLYLRESTDVCITNSFLYAGIV-------------------IN
AgTri      LGVTKLAGLIPSVQARRLHRIAQSSDETMKEFLEKDQMEQLYRKLWVQAGGEREEIPSIW
GaGa       LGLPRLAKAIPEAQVRRLIRVATSSDEVTRKVSYACGISDEVERLWLARGGDMSSVPRFE
SPun       LGIPRLERLIPSVQARSLHRISQSSDYKIMNIAFSCGLESEFEKKWVQAGGAKETRPNLK
PlaMe      LGVTRLASLIPSIQARRLHRIATSEDETIRNIAMANNIEEEFQNLWVTAGGKKEEIPRIT
PBla       LGITRLETTLPHQRCNTLLKIINSPDLTTRTIARALGIEEQFSKWWKKAGGSEDNKPKIY
           *.:. :   :   : . :  :  .             :

LaMa       -----PCRVRNEIV---------------------------CSSKEAQVEWEKIWRTSAD
NePa       -----P---TLPIF---------------------------ESKSAIKQHWADQLLQSID
CroHe      -----PCRLNSTVV---------------------------TSKTEVREEWGNMLSNSVD
DeFu       DVIDEIIKTHH------LHILHA-ECLNMSLELTASY----NWRESEFNRWKDM---VVQ
AgTri      E----ALPTSEQSNRG--DTTSEWEAPILKPKFPKPC----DWRKKEFEKWTKL---VSQ
GaGa       D----PEAPRSPGVQGPCEAAQEIPSVVRKLAIPRPS----NWRAEEHSKWAQL---SCQ
SPun       DILPPPPEPKY------YEKLNEWEKPKLKIKFPLPL----NYRKEEFREWAKL---KCQ
PlaMe      D----PVSIDYRLPRRILELLNEWEKPAPKKMYPIPC----NWREAEMAHWKNL---PCQ
PBla       P-----------------QPQNTTQKPTEKEEIPVTTKIQPTHKKKEAEEWAKL---PCQ
                                                            .* .      :

LaMa       GRSGVNEDVDPASYTWVGKPRRVFPRIHLRGIQLRGGTLNTKARASRGRETAPSDIACRG
NePa       GRDLAQTDIDVASHWWIQHPENVFPRLHLRGLQLRGGLLSTKTRNNRGPTRQ-ANVSCVG
CroHe      GRELRHPEVDKASHEWLLRPDRVFPRLHMRGIQLRGGLLPTKARSGRGNNRQVTSKTCRG
DeFu       GDGVQYYQNNACSNYWLMKPKNMKENQYIAALKLRANIPDTRETLTRGR-NR-QFSNCRY
AgTri      GRGISSFKNDSISNDWLQYYRRIPHRKLITALQLRANVYPTREFLSRGREEN-STKACRH
GaGa       GEGVELFRNDPVSNGWINGRGRLAERLRIVALKLRSNVYPTREFLGRGM-AG-TNVGCRH
SPun       GSGIQHFQDDIISNNWLRINKGITEKQFILALKLRANVFPTREFLGRGT-QN-ANRACRH
PlaMe      GSGIEHFDNDTISNDWLQFHRGFSERQFLMGLKIRANVYPTREYQGRGR-TN-KNVNCRN
PBla       GGAVENFINDSISNDWIRYKKDMTEKEFITAIKLRTNTYPTREFLARGK-EN-YERKCRK
           *        :  *  *:     .  .  : .:::* .   *:    **         *

LaMa       ACHARETLNHILQICEVTHDARCARHNRIAKLLEKMLRRRVDRTWIEPIIP-TQKSFIKP
NePa       ACRMPESINHILQVCEKTHDARCARHNRVMKKVNQLLRAKGLKSWMEPIIP-TSTTFMKP
CroHe      ACQNVETLNHILQVCDVTHDARCARHNRVLHRLERLLHKKGLRTTLEPVIP-AGSSFIKP
DeFu       CMDVGETISHISGSCPKTKDIRIRRHNKILELLIERAKRCGWIAYREPHLRNPNGELRKP
AgTri      CNVENETCAHIIGQCPVTKDARIKRHNHICDMLSEEARKKDWVVFKEPYIRDTTKELYKP
GaGa       CTHPRETLGHILGICPSVQEARILRHNKLCKILAAEGKKCEWTVYYEPHLRNAAGELRKP
SPun       CKAKFETCSHILGQCPTLQEARIKRHNKLCTLLKEEAKKLEWEVFDEPHLHNRAGELRKP
PlaMe      CTASYESLSHILGQCPAVQGARIRRHNKLCSMLKREAKELKWVVYEEPHLHTTEKELRKP
PBla       CPAKYESLSHILGQCPGLQNARIWRHNAIGKLLANQAKKKKWTVHQEPHLRNKDNQLRKP
           .    *:  **   *   :  *  *** :   :    :        ** :      : **

LaMa       DLLVDTGKRIIILDVTVVAEPR---MEQSHQLKSSKYGSPDNTAAIVAWTG--TKVPIKH
NePa       DILVESAGKLIVMDVSIVAGHR---LPETWNLKTSKYGSNQATQDLQRWHN--SLISIKH
CroHe      DLIIETDTQTYVLDVSVVAGYR---MTESWDIKVEKYGRPDRVEAIRKWAAIPATKRVEH
DeFu       DLILTKNNTAHVVDVSIRYETSAGSLESAFEQKVSYYS--ELMSAIQQHTK---CSQVAF
AgTri      DLIFVKDGHAFVVDVTVRYETTTTSLEEAAAEKVNKYK--HLETEVRNLTN---AKDVVF
GaGa       DLIFVRDGTALVVDVTVRYEKDSASLSAAAADKAAKYL--GLNAQIQELTG---AEHVTY
SPun       DVILVKDKKALVVDVTVRFEYKEDGLRKAAKEKASYYS--DLTEPIIELTK---AKKVNY
PlaMe      DLIFVKEEMALVVDVTVRFEYKEKVFEDAAAEKVRHYK--DLTSQIKELTG---AKEIEY
PBla       DYIFTKERTAMVVDITVRWEGKMGGLEEAAQEKIKYYE--ELRDTIQKEWG---VKDISF
           * :.       ::*:::        :  :   *   *        :           : .

LaMa       VPVVLSSRGLLYGPSGRGLRTMGLTSRDLCD----LCLLAVAGSLKCYDAYTRGT-----
NePa       VPVIISARGLLFNKSGQGLRDLGLTRRDISD----ICLLSIIGSLKSYDLYKRGTLDDFR
CroHe      LPVIVSNRGLCYQPSGDGLRALGLTSRDISD----LCLLAISGSLRCYDLYMRGTRAV--
DeFu       HGFIVGCRGTWPTCNNNLLDKLGIQYPNPFK--KNICFTTLMDTLKMFQIFMDN------
AgTri      MGFPLGARGKWYTGNFELLNTLGLSKSRQVRVAKTLSTVALLSSVDIVHMFASRARRVQ-
GaGa       FGFPIGARGKWHADNGEVLSELGLSRSRKERVARLLSWRALLGSVDMVNIFASKHRQETL
SPun       FGLPIGARGKWPPENNNLLRELGMEEGRVKRLARLFSLRTLLYSTNMLHMFENIGKGQKA
PlaMe      FGFPLGARGKWPEINEKVLTALGMPDYQQKRTAKRFSKRTLLYSIDVINTFENIGKNNK-
PBla       HGFVCGARGKWPDQNFGILTKLGIESHRLDGLAKLFSRRTVQYSCRMLHTFERDKKNQTG
             .  . **     .   *  :*:           :.  ::  :    . :

LaMa       ------------
NePa       RTKPR---SGIG
CroHe      ------------
DeFu       ------------
AgTri      --------DSG-
GaGa       LDDAPACVEHVA
SPun       Y-----FVNSNL
PlaMe      --------NNVP
PBla       HVDC--------
```

**Figure S11:** MSA R2 proteins selected for activity testing: LaMa (*L. marequensis*, Largescale yellowfish), NePa (*N. papilliferus*, killifish), DeFu (*D. fucus*, Northern dusky salamander), CroHe (*C. o. helleri*, Southern Pacific Rattlesnake), PBla (*P. blainvillii,* Blainville's Horned Lizard), SPun (*S. punctatus*, Tuatara), GaGa (*G. gangeticus*, Gharial crocodile), AgTri (*A. tricolor*, Tricoloured blackbird) and PlaMe (*P. megacephalum*, Big-headed turtle). Myb2 domain highlighted in orange. N-terminal ZnFs annotated with red asterisk.

#### Supplementary Methods

##### Supplementary Figure 12

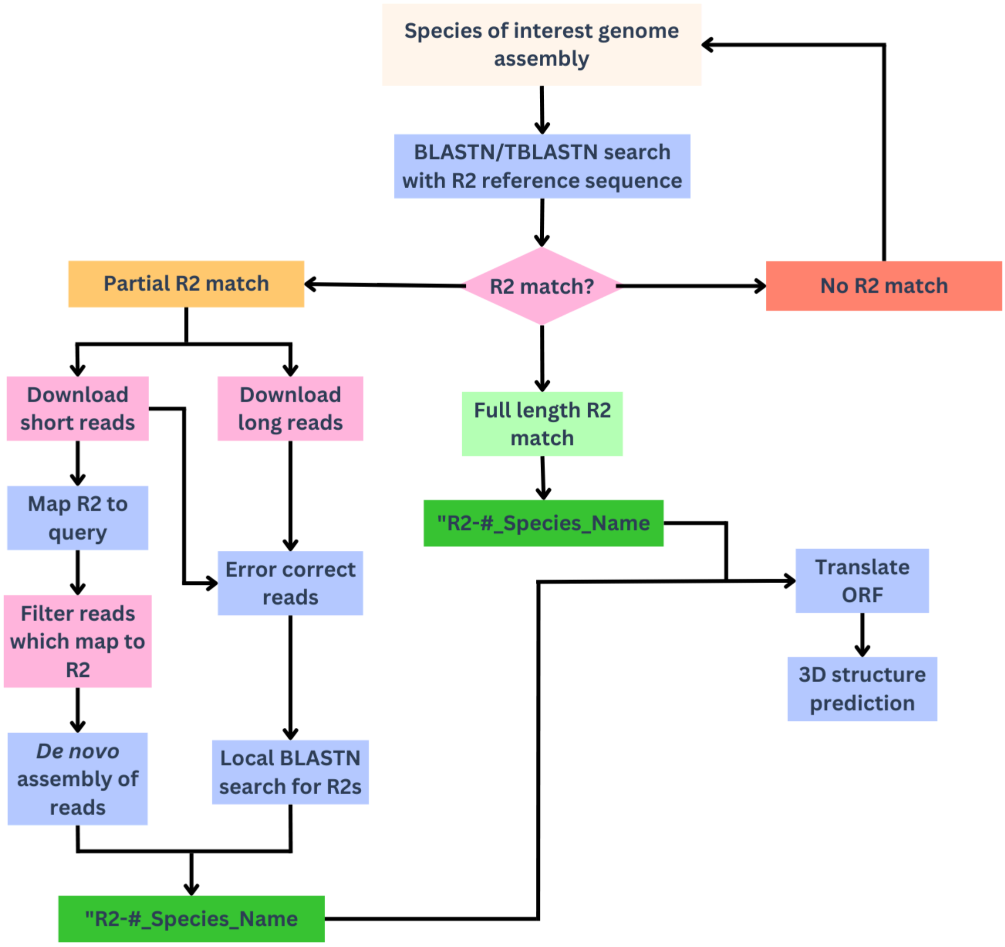


Figure S12: Overview of the R2 discovery pipeline. A sensitive online BLASTN and TBLASTN search was used to find the presence of R2s in the species of interest. Based on the outcome of the search, the R2 was either categorised as full-length or flagged for follow-up using the truncated/misassembled pipeline. Species were removed from further analysis if there is no significant match to R2s. For a truncated or misassembled R2, a subset of long reads and short reads were downloaded from the SRA database. Long reads were made into a BLASTN database to find R2-containing reads. R2-containing reads were error-corrected with short reads using Ratatosk. The error-corrected R2-containing long reads were made into a BLASTN database. Using a subset of R2 query sequences, a sensitive local BLASTN search was performed. Sequences that exceed 4200 nucleotides and contain the required 28s rRNA flanking target sites were selected for ORF translation. Finally, the R2 sequences that fulfil all the requirements (28S rRNA flanks, ORFs, domains) were categorised as full-length R2s.

**Downloading long reads from SRA database and sensitive BLASTN search**

```
#!/bin/bash
#configure to your HPC requirements
#Required tools
#BLAST+, SRA-Toolkit, seqtk, fastqc

#Download long reads
wget https://sra-LONG-READS
#Convert SRR to fastq
fastq-dump SRR-LONG-READS

#Convert fastq to fasta
seqtk seq -a lONG-READS.fastq > lONG-READS.fasta

#Run fastqc on data
#fastqc lONG-READS.fasta
#Concatanate fasta if there are multiple
#cat *lONG-READS.fasta  > LONG-READS-COMBINED.fasta

#Make blast database using long reads
makeblastdb -in lONG-READS.fasta -dbtype nucl -out LONG-READS_db
#Sensitive blast search using R2 query sequences
blastn -query R2-QuerySeqs.fasta -db LONG-READS_db -word_size 7 -outfmt "6 qseqid
sseqid sseq pident length mismatch gapopen qstart qend sstart send evalue bitscore"
> LONG-READS_best_matches

#Extract best hits as fasta file
awk '{print ">"$2"\n"$3}' LONG-READS_best_matches > seqs.fasta
```

**Downloading short reads from SRA database and de novo assembly of R2s**

```
#!/bin/bash
#configure to your HPC requirements

#Required tools
#BWA, SAMtools, SPAdes, SRA-Toolkit, fastqc

#Download short reads
wget https://sra-SHORT-READS

#Convert SRR to fastq
fastq-dump --split-e SHORT-READS

#Run fastqc on data
#fastqc SHORT-READS_1.fastq SHORT-READS_2.fastq

#Index R2 query sequence
bwa index R2-QuerySeqs.fasta

#Map short read sequences to query file
bwa mem R2-QuerySeqs.fasta SHORT-READS_1.fastq SHORT-READS_2.fastq > aln.sam

#Convert sam to bam
samtools sort -O BAM -o aln.bam aln.sam

#Index bam file
samtools index aln.bam

#Extract mapped reads
samtools view -F 4 -b -o mapped.bam aln.bam

#Convert mapped reads to fastq file
samtools fastq mapped.bam > mapped.fastq

#Make spades directories
mkdir spades_denovo
makdir spades_ts

#De novo assembly of R2 mapped reads with Spades denovo and with trusted contigs
options
spades.py --12 mapped.fastq -o spades_denovo
spades.py --12 mapped.fastq --trusted-contigs R2-QuerySeqs.fasta -o spades_ts
```

**Downloading long and short reads from SRA database and error correction prior sensitive BLASTN search**

```
#!/bin/bash
#configure to your HPC requirements

#Required tools
#BLAST+, BWA, SAMtools, SPAdes, SRA-Toolkit, fastqc

#Download short reads
wget https://sra-SHORT-READS

#Download long reads
wget https://sra-long-READS

#Convert SRR to fastq
fastq-dump --split-e SHORT-READS
fastq-dump SRR-LONG-READS

#Run fastqc on data
fastqc SHORT-READS_1.fastq SHORT-READS_2.fastq
fastqc LONG-READS.fasta

#Concatanate long reads
cat *.fastq > LONG-READS.fastq

#Create a list file containing short read headers named short_reads.lst

#Error correct long reads with short reads with Ratatosk
Ratatosk correct -v -c 32 -s short_reads.lst -l LONG-READS.fastq -o
corrected_LONG-READS

#Convert fastq to fasta
seqtk seq -a lONG-READS.fastq > lONG-READS.fasta

#Make blast database using error-corrected long reads
makeblastdb -in lONG-READS.fasta -dbtype nucl -out LONG-READS_db
#Sensitive blast search using R2 query sequences
blastn -query R2-QuerySeqs.fasta -db LONG-READS_db -word_size 7 -outfmt "6 qseqid
sseqid sseq pident length mismatch gapopen qstart qend sstart send evalue bitscore"
> LONG-READS_best_matches

#Extract best hits as fasta file
awk '{print ">"$2"\n"$3}' LONG-READS_best_matches > seqs.fasta
```

**Plotting the evolutionary rate of individual residues**

```
# Load libraries
library(ggplot2)
#Create MSA of aligned RT sequences
#Use IQTree to run site-specific algorithm
http://www.iqtree.org/doc/Advanced-Tutorial
#Extract the output file with site rates mapped to individual positions in the MSA

# Define sequence
sequence <- unlist(strsplit("YOUR-RT-SEQ-OF-INTEREST-FROM-MSA", NULL))

# Read site rates from the CSV file
numbers <- read.csv("~/site-rates.csv")

# Set this to TRUE if you want to filter out dashes in MSA, or FALSE if you want to
keep
filter_dashes <- TRUE

# Create a data frame mapping positions, residues, and numbers
mapping <- data.frame(
  Position = seq_along(sequence),
  Residue = sequence,
  Number = numbers$Number
)

# Apply the dash filter conditionally
if (filter_dashes) {
  mapping <- mapping %>%
    filter(Residue != "-") %>%
    mutate(Position = seq_along(Residue))
}

# Split residue plot into three rows
mapping <- mapping %>%
  mutate(Group = ceiling(seq_along(Residue) / (n() / 3)))

# Custom colours
subdued_palette <- c("#2F7AC6", "#4D9DC6", "#7EC0B8", "#A7D2A0", "#D1E0A8",
"#F2F4B0")

# Plot using ggplot2
ggplot(mapping, aes(x = Position, y = 1, fill = Number)) +
  geom_tile() +
  geom_text(aes(label = Residue), family = "Courier", size = 3, color = "black") +
  scale_fill_gradientn(colors = subdued_palette) +  # Use custom subdued gradient
  scale_x_continuous(breaks = mapping$Position, labels = mapping$Residue) +
  theme_minimal() +
  theme(
    axis.title.y = element_blank(),
```

```
    axis.text.y = element_blank(),
    axis.ticks.y = element_blank(),
    panel.grid = element_blank(),
    axis.text.x = element_text(family = "Courier", size = 5, vjust = 1),
    axis.ticks.x = element_line(),
    axis.title.x = element_text(vjust = -0.5)
  ) +
  facet_wrap(~ Group, nrow = 3, scales = "free_x") +
  labs(title = "Protein Sequence Mapping", x = "Residue", fill = "Number")

ggsave(file = "~/sites.svg")

print(mapping)
```

**Plotting PSSM charts**

```
# Load libraries
if (!requireNamespace("ggrepel", quietly = TRUE) | !requireNamespace("ggforce",
quietly = TRUE)) {
  install.packages(c("ggrepel", "ggforce"))
}
library(dplyr)
library(ggplot2)
library(ggrepel)
library(ggforce)

#Run PSSM using PSIblast for RTs of interest using 1) BoMo then 2) TaGu RTs as the
query
#Create csv: R2 IDs, corresponding bitscores, common species group, number of
N-terminal ZnFs
#For example:
#ID_Code, BoMo, TaGu, ZnF_No, Common.Group
#R2-1_OL, 150, 400, 3, Fish (Bony)

# Load data
df <- read.csv("~/PSSMscores.csv")
str(df)

# Validate columns
required_cols <- c("BoMo", "TaGu", "Common.Group", "ID_Code", "ZnF_No")
if (!all(required_cols %in% colnames(df))) stop("Required columns are missing.")

# Prepare the data
df_filtered <- df %>%
  select( BoMo = BoMo,  TaGu = TaGu, ZnF_No, Common.Group, ID_Code) %>%
#Enter common group you want to plot, e.g. “Fish (Bony)”
  filter(Common.Group %in% c("Common.Group of interest"))
if (nrow(df_filtered) == 0) stop("The filtered data frame is empty.")

#Filter any R2s from the plot if needed
#Filter the data frame to remove rows where 'label' is in 'labels_to_remove'
df_filtered <- df_filtered %>%
  filter(!Code %in% c("R2-1_Example"))

# Variables for axes
x_var <- "TaGu"
y_var <- "BoMo"
# Create the PSSM plot
ggplot(df_filtered, aes_string(x = x_var, y = y_var, color = "Common.Group")) +
  geom_point(show.legend = FALSE) +  # Remove legend for points
  #geom_point(position = position_jitter(width = 0.2, height = 0.2), show.legend =
FALSE) +  # Remove legend for jittered points
  # Draw hulls based on ZnF_No, ignoring Common.Group: this groups and colour codes
R2s based on the number of N-terminal zinc fingers and not common species group
```

```
  geom_mark_hull(aes(group = ZnF_No, fill = ZnF_No),
                 concavity = 10, expand = unit(2.5, "mm"),
                 alpha = 0.15, colour = "black", size = 0.0, show.legend = FALSE) +
# Remove legend for hulls
# Colour code for each species group
  labs(x = x_var, y = y_var) +  # No legend for color and fill
  geom_text_repel(aes(label = Code), size = 3) +
  theme_minimal() +
  theme(
    panel.grid.major = element_line(),
    panel.grid.minor = element_line(),
    axis.line = element_line(color = "black", size = 0.3)  # Keep axis lines
  ) +
  scale_color_manual(values = c(
    "Fish (Bony)" = "#76B49E", "Tunicata" = "#6495EC", "Arthropoda" = "#B196FF",
    "R/Bird" = "#6B8E22", "Echinodermata" = "#4169E1", "R/Snake" = "#FFBF00",
    "R/Lizard" = "#E97929", "R/Crocodile/Alligator" = "#9ACD31", "Cnidaria" =
"#5661D6",
    "Ctenophora" = "#271F5B", "Fish (Cartilaginous)" = "#87CEEB", "R/Turtle" =
"#E0E41D",
    "Amphibian" = "#008080", "Mollusca" = "#B65090", "Annelida" = "#E08BE0",
    "Platyhelminthes" = "#DFB0FF", "R/Lizard (Like)" = "#AE3E0D", "Porifera" = "red"
  )) +
  scale_fill_manual(values = c("A" = "#0B8731", "D" = "#2B10FF", "B" = "#FC8D62",
"C" = "#FC8D62"))

ggsave(filename = "fishesPSSM.svg", width = 10, height = 10, path = "~/your-path")
```

**Supplementary Figure 13**

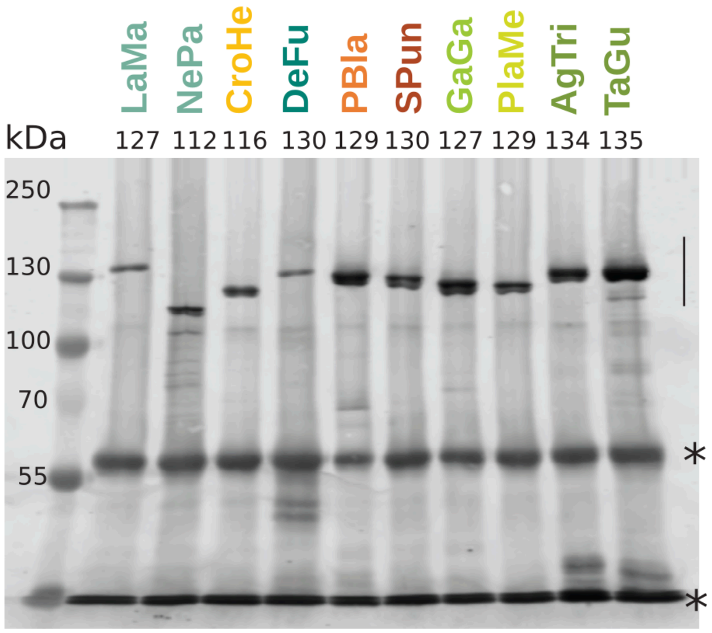


**Figure S13:** Related to Fig. 4b-c. Immunoblot of affinity purified R2 proteins resolved from FLAG affinity by 8% SDS-PAGE. Asterisks denote background bands from flag antibody purification beads. LaMa (*L. marequensis*, Largescale yellowfish), NePa (*N. papilliferus*, killifish), CroHe (*C. o. helleri*, Southern Pacific Rattlesnake), DeFu (*D. fucus*, Northern dusky salamander), PBla (*P. blainvillii*, Blainville's Horned Lizard), SPun (*S. punctatus*, Tuatara), GaGa (*G. gangeticus*, Gharial crocodile), and PlaMe (*P. megacephalum*, Big-headed turtle), AgTri (*A. tricolor*, Tricolored blackbird), and TaGu (*T. guttata*, Zebra finch)

#### Supplementary Figure 14

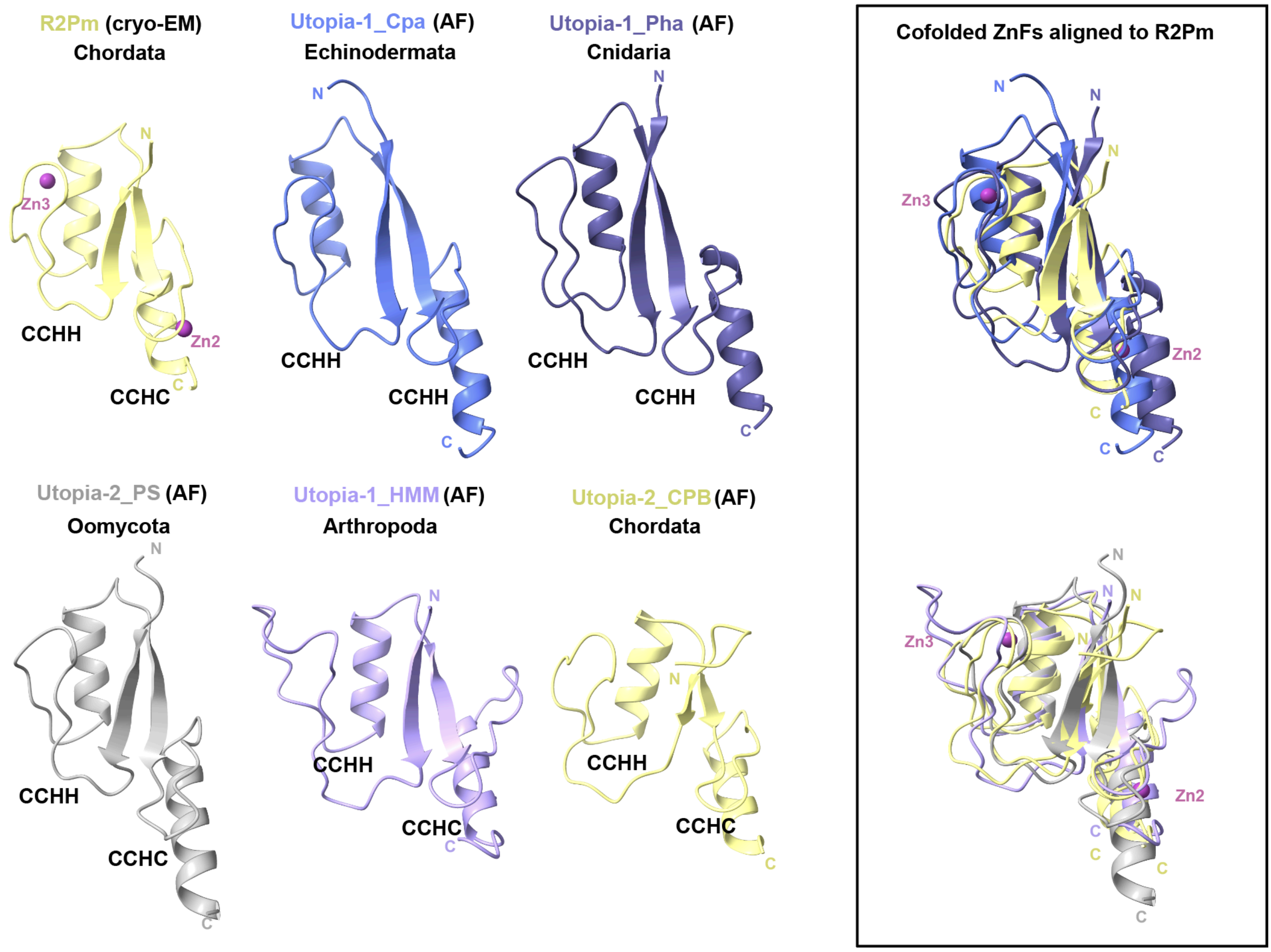


**Figure S14:** Cofolding ZnFs are a feature shared among R2 A-lineage and expected for some Utopia. Protein segments that encode for cofolding ZnFs are displayed, which correspond to ZnF3:2 in the experimental structure of PlaMe (R2Pm) and ZnF2:1 of all Utopia elements indicated (AlphaFold models, AF). Individual protein folds are coloured according to the phyla discussed in the main text. Structural alignments for each row relative to R2Pm ZnF3:2 fold are made on the right (boxed), where the top panel includes three structures and the bottom alignment a total of four. The names of the host species not provided in the main text include those annotated by us, *Crossaster papposus* (Cpa), *Phenganax stokvisi* (Pha), and those previously annotated (4) *Phytophthora sojae* (PS), *Heliconius melpomene* (HMM), *Chrysemys picta bellii* (CPB).
